# Personalized whole-brain activity patterns predict human corticospinal tract activation in real-time

**DOI:** 10.1101/2024.08.15.607985

**Authors:** Uttara U Khatri, Kristen Pulliam, Muskan Manesiya, Melanie Vieyra Cortez, José del R. Millán, Sara J Hussain

## Abstract

**BACKGROUND:** Transcranial magnetic stimulation (TMS) interventions could feasibly treat stroke-related motor impairments, but their effects are highly variable. Brain state-dependent TMS approaches are a promising solution to this problem, but inter-individual variation in lesion location and oscillatory dynamics can make translating them to the poststroke brain challenging. Personalized brain state-dependent approaches specifically designed to address these challenges are therefore needed.

**METHODS:** As a first step towards this goal, we tested a novel machine learning-based EEG-TMS system that identifies personalized brain activity patterns reflecting strong and weak corticospinal tract (CST) output (strong and weak CST states) in healthy adults in real-time. Participants completed a single-session study that included the acquisition of a TMS-EEG-EMG training dataset, personalized classifier training, and real-time EEG-informed single pulse TMS during classifier-predicted personalized CST states.

**RESULTS:** MEP amplitudes elicited in real-time during personalized strong CST states were significantly larger than those elicited during personalized weak and random CST states. MEP amplitudes elicited in real-time during personalized strong CST states were also significantly less variable than those elicited during personalized weak CST states. Personalized CST states lasted for ∼1-2 seconds at a time and ∼1 second elapsed between consecutive similar states. Individual participants exhibited unique differences in spectro-spatial EEG patterns between personalized strong and weak CST states.

**CONCLUSION:** Our results show for the first time that personalized whole-brain EEG activity patterns predict CST activation in real-time in healthy humans. These findings represent a pivotal step towards using personalized brain state-dependent TMS interventions to promote poststroke CST function.

## Introduction

Transcranial magnetic stimulation (TMS) is a noninvasive brain stimulation technique that could feasibly treat a variety of psychiatric and neurological disorders, including depression (George et al., 1995, 2010) obsessive-compulsive disorder (Mantovani et al., 2006; Tendler et al., 2021), memory deficits (Freedberg et al., 2022; Solé-Padullés et al., 2006; Wang et al., 2014), cognitive decline (Luber & Lisanby, 2014), and motor impairments caused by neurological damage (Bunday & Perez, 2012; Di Lazzaro et al., 2008; Du et al., 2016; Jo & Perez, 2020). Early studies showed that TMS interventions delivered to the sensorimotor cortex can alter corticospinal tract (CST) transmission (Huang et al., 2005; Pascual-Leone et al., 1995, Chen et al., 1998). Given the mechanistic role of the CST in voluntary upper extremity movement (Lemon, 2008) and the prognostic utility of CST integrity in predicting poststroke upper extremity motor recovery (Stinear et al., 2007), TMS interventions that upregulate CST transmission could feasibly improve voluntary motor function in individuals with stroke-related disruption of the CST. However, it has recently become apparent that the effects of TMS interventions on CST transmission are highly variable both within and between individuals (Hamada et al., 2013; López-Alonso et al., 2014), such that conventional TMS interventions do not reliably upregulate CST transmission even in healthy adults.

TMS applied over the sensorimotor cortex trans-synaptically activates CST neurons (Di Lazzaro & Ziemann, 2013; Hoogendam et al., 2010; Mills et al., 1992), resulting in a peripheral muscle response termed a motor-evoked potential (MEP). The peak-to-peak amplitude of an MEP reflects the magnitude of CST activation at the precise moment of TMS delivery. Yet, MEP amplitudes dynamically fluctuate over time, even when keeping other parameters such as stimulation location and intensity constant (Jung et al., 2010; Kiers et al., 1993). Such dynamic fluctuations can be attributed in part to variability in subthreshold depolarization of CST neurons and cortical interneurons synapsing onto them (Di Lazzaro & Ziemann, 2013; Ziemann et al., 1996). Consistent with this notion, accumulating evidence has shown that CST activation depends on ongoing sensorimotor oscillatory activity at the time of stimulation (Berger et al., 2014; Bergmann et al., 2019; Hussain et al., 2019; Ozdemir et al., 2022; Suresh & Hussain, 2023; Wischnewski et al., 2022; Zrenner et al., 2018), including sensorimotor rhythm phase (Bergmann et al., 2019; Wischnewski et al., 2022; Zrenner et al., 2018), sensorimotor rhythm power (Hussain et al., 2022; Madsen et al., 2019), and interactions between them (Hussain et al., 2019b; Ozdemir et al., 2022; Suresh & Hussain, 2023). For example, TMS more strongly activates the CST during sensorimotor mu rhythm trough than peak phases (Bergmann et al., 2019; Wischnewski et al., 2022; Zrenner et al., 2018), and this effect is strongest during periods of high mu rhythm power (Hussain et al., 2019; Ozdemir et al., 2022; Suresh & Hussain, 2023). These studies raise the possibility that delivering TMS interventions during mu trough phases could enhance their efficacy. Indeed, EEG-triggered repetitive TMS interventions delivered during sensorimotor rhythm mu trough phases increase CST transmission, while identical interventions delivered during mu peak phases weakly depress it (Baur et al., 2020; Zrenner et al., 2018). Thus, coupling TMS interventions to brain activity patterns (i.e., brain states) reflecting strong CST activation could potentiate their therapeutic effects in individuals with poststroke motor impairments.

Although several studies have shown that TMS more strongly activates the CST during sensorimotor mu rhythm trough than peak phases, the magnitude of this effect varies across studies (Bergmann et al., 2019; Hussain et al., 2019b; Madsen et al., 2019b; Wischnewski et al., 2022; Zrenner et al., 2018), suggesting that mu phase-dependent variation in CST activation exhibits substantial inter-individual variability even in healthy adults. Furthermore, translating real-time, mu phase-dependent TMS approaches from the healthy to the poststroke brain can be challenging (Hussain et al., 2020). Because stroke survivors are a highly heterogeneous population, each stroke survivor has a unique pattern of motor impairment and recovery-related adaptive plasticity (Delvaux et al., 2003; Grefkes & Ward, 2014; Jones, 2017; Lotze et al., 2012; Luft et al., 2004; C. Stinear, 2010) that could alter sensorimotor rhythm characteristics and their relationship to CST activation. Lesion-related volumetric brain loss in each stroke survivor is also unique, such that the mapping of brain activity to EEG scalp signals varies across stroke survivors (Lopez-Larraz et al., 2017; Park et al., 2016). Personalized brain state-dependent TMS approaches specifically designed to address these issues are therefore needed.

Consistent with this need, we developed and tested a novel machine learning-based EEG-triggered TMS system that identifies and targets personalized whole-brain activity patterns reflecting time windows when TMS either strongly or weakly activates the CST (i.e., personalized strong or weak CST states) in healthy adults. We first acquired a single training dataset for each participant which included EEG and EMG recorded during single-pulse motor cortex (M1) TMS. We then used this dataset to build a personalized classifier that discriminates between EEG activity patterns during which TMS elicited either a large or small MEP. Finally, we tested this personalized classifier by evaluating MEP amplitudes during real-time, EEG-triggered TMS targeting personalized strong, weak, and random CST states. Our results show that this system can accurately identify and target personalized whole-brain EEG activity patterns corresponding to strong and weak CST activation in real-time. These findings represent a key step towards using personalized, machine learning-driven brain state-dependent TMS interventions to promote poststroke CST function and motor recovery.

## Methods

### Data acquisition

#### Participants

21 healthy adults participated in this single-session study, which involved single-pulse transcranial magnetic stimulation (TMS) during 62-channel electroencephalography (EEG) and bipolar EMG recordings from the left first dorsal interosseous (L. FDI) and left abductor pollicis brevis (L. APB) muscles. Of these participants, one was excluded due to EMG signal corruption, and one was excluded due to excessively noisy EEG signals. Thus, our final sample size was N=19 (15 F, 4 M, age = 20.8 ± 0.7 [standard error of the mean; SEM] years). This study was approved by the Institutional Review Board at the University of Texas at Austin, and all participants provided their written informed consent prior to participation.

#### Experimental design

After experimental setup was complete, the TMS stimulation location and intensity were empirically determined for each participant (see *TMS*). Then, participants completed a 5-minute EEG recording while resting quietly with their eyes open. After resting EEG, 6 blocks of 100 single brain state-independent TMS pulses at 120% of resting motor threshold (RMT) were delivered to the scalp motor hotspot for the L. FDI muscle while EEG and EMG were recorded. Resulting EEG and EMG data were used to build a personalized classifier that could discriminate between whole-brain EEG activity patterns during which TMS strongly activated the CST (i.e., strong CST states) or weakly activated it (i.e., weak CST states), as measured via L. FDI motor-evoked potential (MEP) amplitudes. Afterward, this classifier was used to deliver real-time, single-pulse brain state-dependent TMS to the scalp hotspot for the L. FDI during strong and weak CST states at two different stimulation intensities (120% and 110% RMT). For comparison, single-pulse brain state-independent TMS was also applied to the scalp hotspot for the L. FDI at these same intensities (i.e., random CST states). Throughout the experimental session, MEPs were recorded from both the L. FDI and L. APB muscles. See Figure 1A for a visual depiction of the experimental timeline.

**Figure 1.**
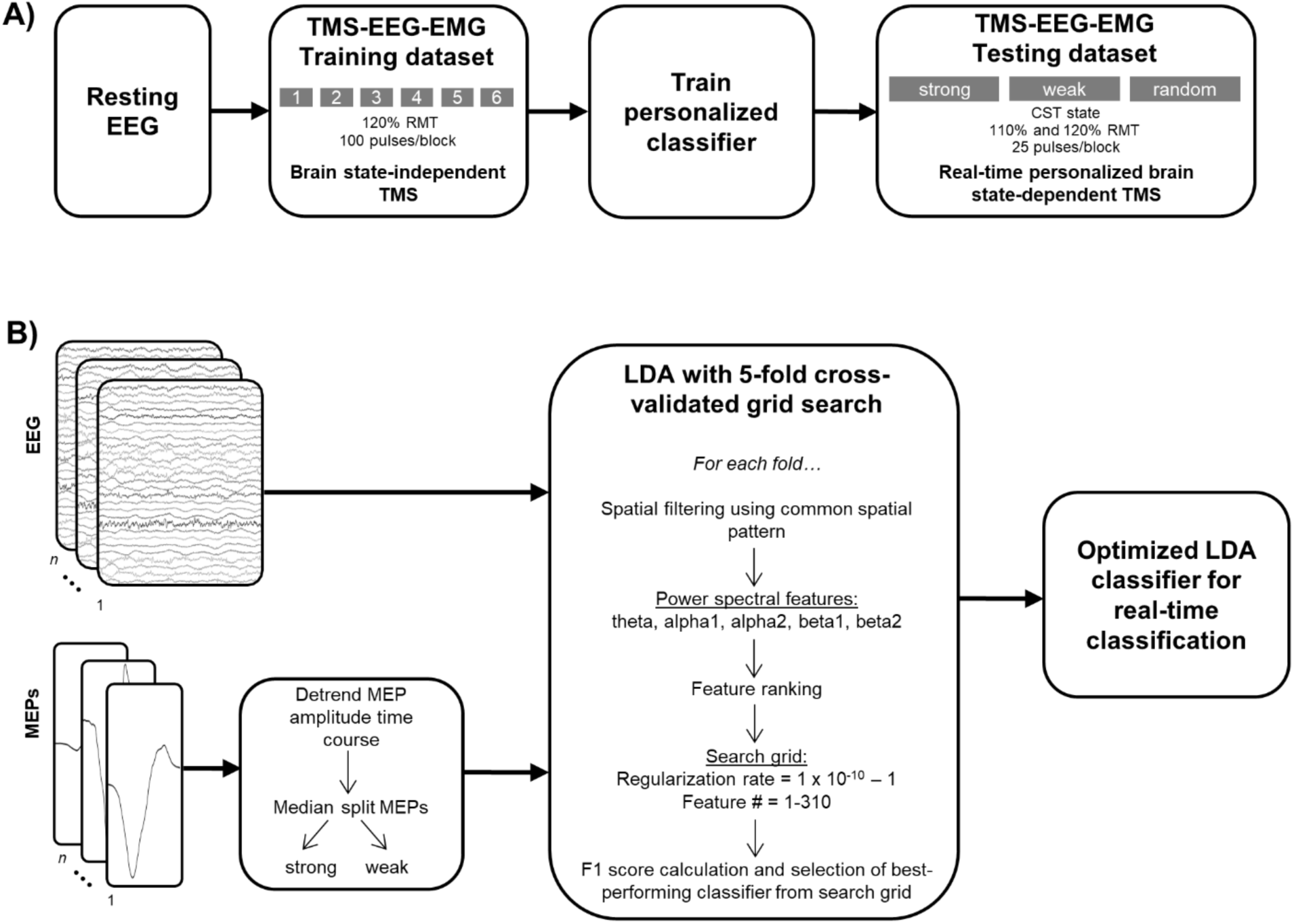
Experimental timeline and machine learning analysis pipeline. A) Experimental timeline. All procedures were completed within a single session. B) Personalized machine learning classifier analysis pipeline.

#### EEG and EMG acquisition

62-channel EEG signals were recorded at 5 kHz (low-pass hardware filtering cutoff frequency: 1250 Hz, 0.001 µV resolution) using TMS-compatible amplifiers (NeurOne Tesla, Bittium Biosignals, Finland). EEG impedances were maintained below 10 kΩ. Bipolar EMG signals were also recorded from the L. FDI and L. APB muscles at 5 kHz (low-pass hardware filtering cutoff frequency: 1250 Hz, 0.001 µV resolution) using Ag-AgCl adhesive electrodes arranged in a belly-tendon montage.

#### TMS

The scalp hotspot was identified over the hand representation area of the right motor cortex as the site at which suprathreshold single-pulse TMS elicited the largest MEPs within the L. FDI as well as a focal muscle twitch. Then, the RMT was determined using a threshold-tracking software tool (MTAT 2.0; Awiszus, 2011). RMT was on average 66 ± 2.3% (range = 51 – 84) of maximum stimulator output. To maximize trans-synaptic activation of corticospinal tract (CST) neurons, TMS was delivered using a figure-of-eight coil held at ∼45 degrees relative to the mid-sagittal line (Mills et al., 1992); Deymed Diagnostic, XT100, biphasic pulse shape). Coil position accuracy was monitored online using frameless neuronavigation (BrainSight, Rogue Research, Inc.).

#### Personalized offline machine learning classification

We acquired a single TMS-EEG-EMG training dataset from each participant by delivering 6 blocks of 100 single TMS pulses to the scalp hotspot for the L. FDI muscle at 120% RMT during EEG and EMG recordings (inter-stimulus interval = 3 s + random jitter). Participants rested quietly with their eyes open during TMS delivery and were provided with short rest breaks between blocks. After acquiring this training dataset, participants rested while a personalized machine learning classifier was built. The purpose of this classifier was to discriminate between whole-brain EEG activity patterns during which TMS either strongly or weakly activated the CST, indexed by L. FDI MEP amplitudes. EEG and L. FDI EMG data were preprocessed using custom-written scripts utilizing the FieldTrip toolbox (Oostenveld et al., 2011) while machine learning classification was performed using custom-written scripts utilizing the MVPA-Light toolbox (Treder, 2020). Both toolboxes operated in the MATLAB environment.

Continuous bipolar EMG data for the L. FDI muscle were divided into segments (-0.100 to +0.400 s relative to each TMS pulse). We then calculated the root-mean-square (RMS) value for each L. FDI EMG pre-stimulus EMG segment (-0.100 to -0.025 s relative to each TMS pulse). Trials contaminated by voluntary muscle activation were identified as those for which pre-stimulus RMS values exceeded a participant-specific threshold, defined as the mean of pre-stimulus L. FDI EMG RMS values + 2 * the standard deviation of pre-stimulus L. FDI EMG RMS values. On average, 2.3 ± 0.4% of all trials (range = 0.2 – 6.8%) were contaminated by voluntary muscle activation per participant. Then, each L. FDI EMG segment was used to calculate peak-to-peak L. FDI MEP amplitudes. To ensure that our classification approach captured trial-by-trial variability in L. FDI MEP amplitudes rather than slow fluctuations in CST activation that can occur with repeated application of single-pulse TMS (Pellicciari et al., 2016), the resulting time course of L. FDI MEP amplitudes was demeaned and linearly detrended, z-transformed, and then rescaled to range between 0 and 1. These steps were performed separately for all L. FDI MEPs not contaminated by voluntary muscle activation per block. Transformed L. FDI MEPs were then combined across blocks and dichotomized into categories that reflected strong activation of the CST (i.e., L. FDI MEPs with amplitudes larger than or equal to the median L. FDI MEP amplitude) or weak activation of the CST (i.e., L. FDI MEPs with amplitudes smaller than the median L. FDI MEP amplitude). These categories were used as class labels during subsequent offline classification.

Continuous 62-channel EEG data were divided into segments (-[0.500 + *x*] to -[0 + *x*] ms before each TMS pulse, where *x* reflects the technical delay of the real-time EEG streaming and analysis system; see *Real-time EEG analysis and personalized brain state-dependent TMS*). To ensure accurate real-time performance, the technical delay was calculated at the beginning of each experimental session and individually adjusted per session. Technical delays were on average 43.6 ± 1.3 (range = 39 - 59) ms. Segmented EEG data were re-referenced to the common average reference, demeaned, linearly detrended, and downsampled to 1 kHz.

For personalized machine learning classification, we used Linear Discriminant Analysis (LDA) with 5-fold stratified cross-validation. As applied here, LDA is a supervised machine learning algorithm that identifies the hyperplane which best separates whole-brain EEG brain activity patterns during which TMS either strongly or weakly activated the CST (i.e., strong or weak CST states, respectively), as indexed by L. FDI MEP amplitudes. We applied a modified version of our previously published personalized classification approach (Hussain et al., 2022) to preprocessed EEG data and L. FDI MEP amplitude class labels.

Trials were randomly divided into folds. For each fold, classifiers were trained on the training dataset (80% of trials) and tested on the testing dataset (20% of trials). We first applied common spatial filter analysis (CSP; Blankertz et al., 2008) to the preprocessed EEG timeseries data. CSP is a signal processing approach that improves the discriminability of two classes of EEG signals by maximizing the variance of EEG data corresponding to one class and minimizing the variance of EEG data corresponding to the other class. CSP as applied here generates subcomponents that reflect spatially filtered EEG timeseries data corresponding to each class. For each fold, CSP spatial filters were calculated using the training dataset and then applied to the testing dataset to avoid information leakage that could bias classification results. All 62 subcomponents generated by CSP were retained and spectrally decomposed using Welch’s method (4-35 Hz with 0.25 Hz resolution). Power spectra obtained for each CSP subcomponent were then summarized by calculating mean spectral power values for each of five canonical frequency bands, including theta (4-8 Hz), alpha1 (8-10 Hz), alpha2 (10-13 Hz), beta1 (13-20 Hz) and beta2 (20-35 Hz). Overall, this approach generated five power spectral features for each of the 62 CSP subcomponents, resulting in a total of 310 power spectral features per participant.

After calculating all features for each fold, we next optimized the number of features included in each participant’s personalized classifier. This was done by first ranking all 310 features in order of the strength of their statistical dependency with L. FDI MEP classes using the chi-square method (Hussain et al., 2022). Here, the negative log of the chi-squared test’s p-value for each feature was taken as its feature score, with higher scores reflecting features that more strongly covary with L. FDI MEP classes. We then used grid search to optimize two aspects of each participant’s classifier: feature number (1 to 310) and regularization rate (100 linearly spaced values from 1× 10^−10^ to 1). During grid search, we iteratively trained multiple classifiers using all possible combinations of feature numbers (with features added in order of importance) and regularization rate values. Overall, this grid search approach produced 31,000 trained classifiers per fold. For each fold, we applied all classifiers trained on the training dataset to the testing dataset. Here, we identified each classifier’s most confident predictions from the testing dataset by calculating the distance of each trial’s prediction from the model’s hyperplane (i.e., each trial’s d-value, with larger d-values indicating more confident predictions). A prediction was labeled confident if the absolute value of its d-value was within the top 50% of that class’s set of d-values. In contrast, a prediction was labeled under-confident if the absolute value of its d-value was in the bottom 50% of that class’s set of d-values. Confident strong CST state predictions (predictions of large L. FDI MEP amplitudes) were labeled 2, confident weak CST state predictions (predictions of small L. FDI MEP amplitudes) were labeled 1, and under-confident predictions were labeled 0. We then calculated each classifier’s prediction performance using the F1 score, focusing only on high confidence predictions. The F1 score is the harmonic mean of precision and recall and was chosen to maximize the number of true positives (i.e., accurate weak CST state predictions) and minimize the number of false negatives and false positives (i.e., inaccurate strong or weak CST state predictions). We then averaged F1 scores across folds and identified the best performing classifier with the fewest features and highest regularization rate. This resulted in selection of one personalized ensemble classifier with 5 LDA models embedded within it (i.e., one per fold). F1 scores were on average 0.68 ± 0.01 when considering only high confidence predictions and 0.63 ± 0.01 when considering all predictions. See Figure 1B for a visual depiction of the personalized classification analysis pipeline.

#### Real-time EEG analysis

After identifying each participant’s best-performing personalized ensemble classifier, we used this classifier to identify EEG activity patterns predicting strong and weak CST activation in each participant in real-time. To achieve this, continuous 62-channel EEG was recorded and streamed to a Dell workstation PC (10 cores, Intel i9 processor, 16 GB RAM, 1 TB Solid State Drive) at 1 kHz using LabStreamingLayer (https://github.com/sccn/labstreaminglayer; Kothe et al., 2024). This workstation was configured to perform real-time EEG analysis in MATLAB version 2020a using a combination of custom-written scripts, FieldTrip, and MVPA-Light Toolboxes.

After being streamed to the workstation PC, EEG data were buffered into overlapping 500 ms windows. Overlap between consecutive windows determined by the technical delay of the real-time EEG streaming and analysis system (with overlap equal to [500ms - technical delay]). Buffered data were downsampled to 1 kHz, re-referenced to common average reference, demeaned, and linearly detrended. Preprocessed data were used to obtain 62 CSP subcomponents per fold using the same CSP parameters used for that participant’s optimized ensemble classifier. This approach produced 5 distinct versions of 62 time-resolved CSP subcomponents (i.e., one per fold). Then, each fold’s CSP subcomponents were used to calculate power spectral features for that fold using the same approach implemented during offline classification. For each fold, the relevant features were selected and used to classify the current EEG segment, resulting in five separate class predictions for each segment. Then, each EEG segment’s predictions were labeled as confident or under-confident using the same procedures applied during offline classification. To obtain a single prediction for each EEG segment, all five predictions were combined using majority voting. If all five predictions were under-confident or a tie occurred, no prediction was made. The label produced by the majority of the five classifiers was chosen as the label for that EEG segment, indicating either a strong CST state, a weak CST state, or no prediction. When 10 consecutive confident predictions of the desired CST state occurred, TMS was triggered. For some participants, 10 consecutive confident strong and/or weak CST states could not be detected in real-time (2 participants for strong states at 120% and 110% RMT, 1 participant for weak states at 110% RMT). In these scenarios, TMS was triggered upon 10 consecutive confident *or* under-confident predictions of the desired CST state.

#### Real-time personalized single-pulse brain state-dependent TMS

TMS was delivered to the L. FDI scalp hotspot and MEPs were recorded from the L. FDI and L. APB muscles. 25 pulses were applied per CST state (strong or weak) and stimulus intensity (110% and 120% RMT) in a blocked manner (minimum interstimulus interval = 3 s + random jitter). In practice, interstimulus intervals were on average 10.0 ± 0.9 and 7.0 ± 0.6 s for strong and weak CST states, respectively. For comparison, we also delivered conventional brain state-independent TMS (i.e., during random CST states) at the same two intensities (interstimulus interval = 3 s + random jitter). Overall, we obtained personalized brain state-dependent MEP amplitudes from 6 blocks of single-pulse TMS for each participant. The order of targeted CST states was counterbalanced across participants. However, stimulation intensities were always tested in the same order, with 120% RMT followed by 110% RMT.

### Data analysis

#### Evaluation of CST state targeting accuracy

After data acquisition was complete, we evaluated the ability of our EEG-triggered TMS system to accurately identify and deliver TMS during pre-defined CST states in real-time. To achieve this, we first divided EEG data obtained during real-time personalized single-pulse brain state-dependent TMS into segments (-[0.500 + *x*] to -[0 + *x*] ms before each TMS pulse, where *x* reflects the session-specific technical delay of the real-time EEG streaming and analysis system). We then applied the same EEG preprocessing, CSP, spectral decomposition, and classification procedures used when performing real-time EEG analysis to these segments, thus mimicking our real-time CST state prediction analysis procedure in an offline analysis environment. For each participant, this approach resulted in a series of predictions made by the offline application of each participant’s personalized ensemble classifier per CST state and stimulation intensity. Then, EEG segments for which the prediction made by the real-time and offline application of each participant’s personalized ensemble classifier were identical were labeled as accurate. For each participant, the percentage of accurate CST states targeted per state and intensity was calculated.

#### Analysis of MEP amplitudes and variability

Continuous L. FDI and L. APB EMG data obtained during real-time, personalized single-pulse brain state-dependent TMS were divided into segments (-0.100 to +0.400 s relative to each TMS pulse). Pre-stimulus EMG data (-0.100 to - 0.025 relative to each TMS pulse) obtained from each muscle were demeaned and linearly detrended. Then, a discrete Fourier transform-based filter was used to attenuate line noise and its harmonics within these pre-stimulus signals. For each muscle, the RMS value for each trial’s processed pre-stimulus EMG signal was calculated. Then, peak-to-peak MEP amplitudes were calculated for each muscle. Trials for which peak-to-peak MEP amplitudes could not be reliably calculated were excluded from analysis (average = 7.2 ± 3.3% of all trials, range = 0 – 43.3%). For each participant, trial-by-trial MEP amplitudes were normalized to the mean of all MEP amplitudes elicited from that muscle at that stimulation intensity. We also evaluated MEP amplitude variability by calculating the coefficient of variation of all remaining MEP amplitudes per CST state, intensity, and muscle for each participant.

#### Characterization of CST state duration

To characterize the mean duration of personalized strong and weak CST states, we applied each participant’s personalized ensemble classifier to their resting EEG data obtained at the beginning of the experimental session. We selected the first 3 minutes of each participant’s resting EEG data and divided these data into consecutive, 500 ms overlapping segments (overlap = 50 ms). EEG segments were then analyzed using the same preprocessing, CSP, spectral decomposition, and classification procedures applied during real-time EEG analysis. We then calculated the average duration of each CST state, including strong CST states, weak CST states, and under-confident states. We also calculated the average time between consecutive similar states (i.e., the inter-state interval) and the percentage of time that each CST state was present.

#### Performance of non-personalized classifiers

After data acquisition was complete, we evaluated the performance of a single, general classifier trained using data from TMS-EEG-EMG training datasets combined across all participants using similar procedures as that described for personalized classification (see *Personalized offline machine learning classification* above). To create training datasets, TMS-EEG-EMG data were compiled across N-1 participants (N = sample size of 19). Testing datasets contained TMS-EEG-EMG data from the remaining held out participant. That is, we performed k-fold cross-validated grid search with k equal to N-1. Given that each fold’s testing set represented an individual participant, F1 performance values were calculated for each fold and then compiled across participants.

#### Pre-stimulus spectro-spatial EEG patterns for personalized strong and weak CST states

We also characterized differences in pre-stimulus EEG brain activity patterns present during classifier-predicted strong versus weak CST states using 500 ms pre-stimulus EEG segments (-[0.500 + *x*] to -[0 + *x*] ms before each TMS pulse, where *x* reflects the session-specific technical delay of the real-time EEG streaming and analysis system). All analyses were performed at the individual participant level. After segmenting EEG data, the same EEG preprocessing procedures used during real-time EEG analysis were applied. All channels of preprocessed EEG data were then spectrally decomposed using Welch’s method using the same parameters used during real-time EEG analysis. The resulting power spectra at each channel were natural log-transformed and averaged across each CST state. The difference between power spectra obtained during strong and weak CST states at each channel was then calculated, and these differences were binned by 1 Hz. This procedure generated a participant-specific matrix of power spectral differences at each channel and frequency. To obtain a group-level representation of differences in power spectra at each channel between CST states, we averaged these matrices across participants.

### Statistical analysis

All statistical analyses were performed using RStudio. Alpha was equal to 0.05 for all comparisons, and all data are expressed as mean ± SEM.

#### CST state targeting accuracy

For each state and stimulation intensity, the percentage of trials during which real-time, personalized brain state-dependent TMS accurately targeted the desired CST state were compiled across participants and compared to theoretical chance (0.5) using separate single-sample, right-tailed Wilcoxon signed-rank tests after confirming deviations from normality using the Shapiro-Wilk test.

#### MEP amplitudes

We evaluated real-time personalized brain state-dependent variation in MEP amplitudes regardless of the intensity used (110% and 120% RMT) or muscle from which they were recorded (L. FDI and L. APB) using a trial-by-trial linear mixed-effects model. This model included natural-log transformed MEP amplitudes as the response variable, STATE, INTENSITY, MUSCLE, and their two- and three-way interactions as fixed effects, pre-stimulus background RMS EMG and inter-stimulus interval as continuous covariates, and PARTICIPANT as random intercept. Model fits were visually inspected using histograms of residuals and quantile-quantile plots. The significance of fixed effects was determined using likelihood ratio tests. Significant fixed effects were evaluated further using pairwise post hoc comparisons. All post hoc comparisons were adjusted for multiple comparisons using the False Discovery Rate correction (Benjamini & Hochberg, 1995). Given that we trained personalized classifiers using L. FDI amplitudes elicited at 120% RMT, we also performed planned pairwise comparisons between CST states for this muscle and intensity.

#### MEP variability

A linear mixed-effects model was also used to evaluate real-time personalized brain state-dependent MEP amplitude variability for both intensities (110% and 120% RMT) and muscles (L. FDI and L. APB). This model included natural-log transformed MEP amplitude coefficients of variation as the response variable, STATE, INTENSITY, MUSCLE, and their two- and three-way interactions as fixed effects, trial-averaged pre-stimulus background RMS EMG and trial-averaged inter-stimulus intervals as continuous covariates, and PARTICIPANT as the random intercept. Model fits were visually inspected using histograms of residuals and quantile-quantile plots. The significance of fixed effects was determined using likelihood ratio tests. Significant fixed effects were further characterized via pairwise post hoc comparisons using the False Discovery Rate correction.

#### Temporal characteristics of CST states

The mean duration of each state, the proportion of time spent in each state, and the time between consecutive similar states were each compared across CST states using separate Kruskal-Wallis rank sum tests for each response variable after confirming deviations from normality using the Shapiro-Wilk test. Pairwise post hoc comparisons were performed using two-sample, two-tailed Wilcoxon Signed Rank tests followed by correction for multiple comparisons using the False Discovery Rate correction.

#### Comparing personalized and non-personalized classifier performance

F1 scores indicating personalized and non-personalized classifier performance were compared using two-sample, paired Wilcoxon Signed Rank tests after confirming deviations from normality using the Shapiro-Wilk test.

#### Relationships between personalized classifier performance and brain state-dependent variation in MEP amplitudes

For each participant, we calculated the percentage difference in MEP amplitudes elicited during strong versus weak CST states and strong versus random CST states at the same muscle and stimulation intensity used during classifier training (L. FDI MEP amplitudes at 120% RMT). These percentage difference values were then regressed against F1 scores obtained from each participant’s personalized classifier using separate Spearman’s correlations after confirming deviations from normality using the Shapiro-Wilk test.

## Results

### CST state targeting accuracy

We first examined the ability of our machine learning-based EEG-triggered TMS system to identify personalized CST states in real-time by calculating the percentage of all trials during which it accurately targeted the desired CST state at each stimulation intensity (Figure 2). Targeting accuracy significantly exceeded chance for strong and weak CST states at 120% and 110% RMT (p < 0.001 for all). For 120% RMT, targeting accuracy was on average 94.5 ± 1.1% and 89.5 ± 2.7% for strong and weak CST states, respectively. For 110% RMT, targeting accuracy was on average 95.1 ± 1.8% and 83.1 ± 6.3% for strong and weak CST states, respectively.

**Figure 2.**
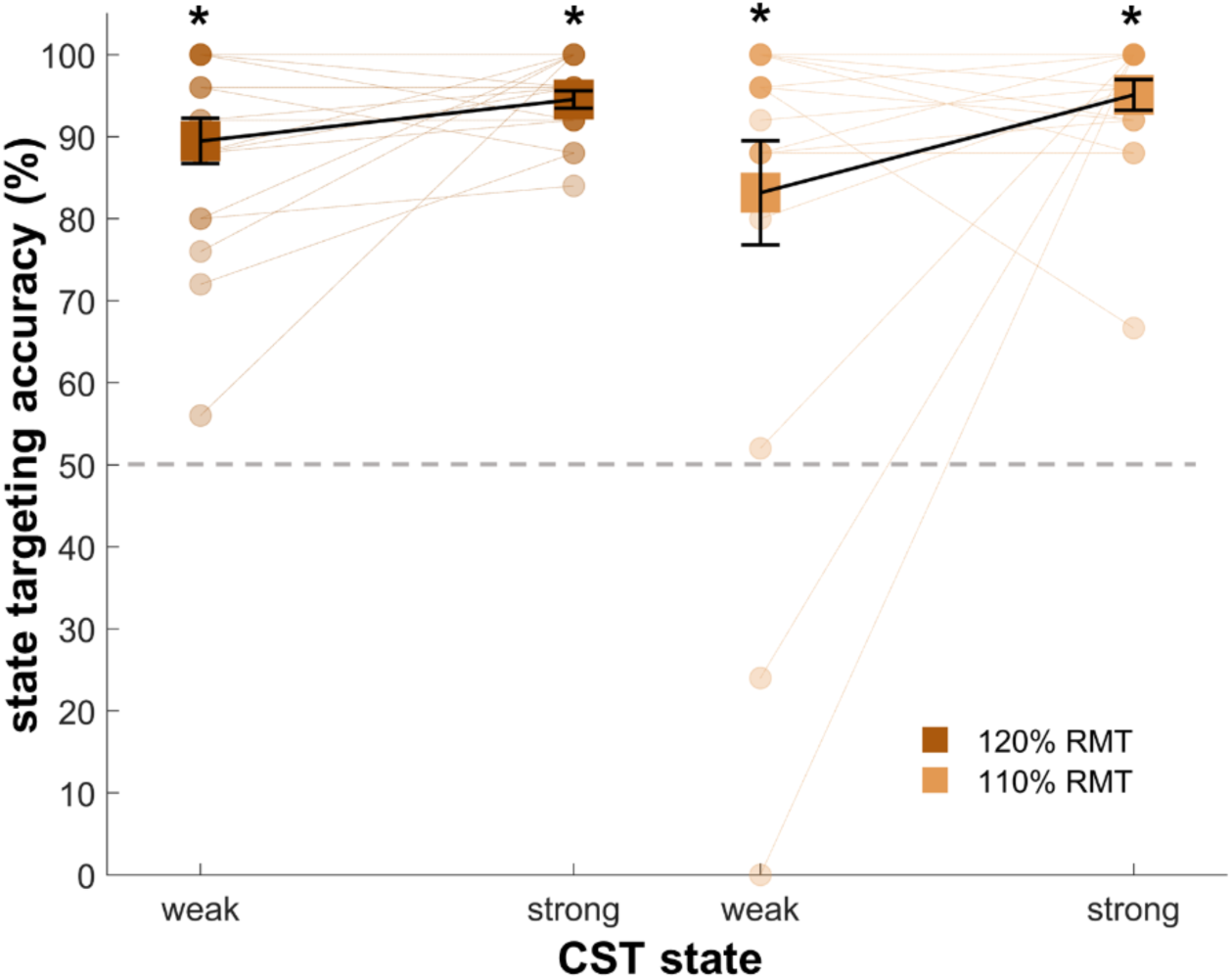
Real-time personalized CST state targeting accuracy. Targeting accuracy of real-time EEG-triggered TMS for personalized strong and weak CST states at 120% and 110% RMT. Asterisks reflect statistically significant comparisons between targeting accuracy and chance level for each combination of state and stimulation intensity. Squares denote group averages, dots denote data from individual participants, error bars denote SEM, and the dashed horizontal grey line denote the theoretical chance level (50%).

### MEP amplitudes

After confirming accurate real-time targeting of personalized CST states, we next evaluated differences in MEP amplitudes elicited in real-time during personalized strong, weak, and random CST states regardless of the intensity used (110% and 120% RMT, Figure 3) or the muscle from which they were recorded (L. FDI and L. APB). A linear mixed-effects model revealed a significant main effect of STATE (likelihood ratio test: F = 11.7, p < 0.001), INTENSITY (likelihood ratio test: F = 28.1, p < 0.001), and MUSCLE (likelihood ratio test: F = 25.4, p < 0.001), as well as a significant two-way interaction between STATE and INTENSITY (likelihood ratio test: F = 5.7, p = 0.003). Post hoc pairwise comparisons revealed that at 120% RMT, MEPs elicited in real-time during personalized strong CST states were significantly larger than those elicited in real-time during personalized weak (p = 0.036) and random CST states (p < 0.001). At 120% RMT, MEPs elicited during personalized weak CST states were also significantly larger than those elicited during random CST states (p = 0.001). At 110% RMT, MEPs did not differ between personalized strong, weak, or random CST states (p > 0.33 for all). Planned comparisons for L. FDI MEPs elicited at 120% RMT showed that MEP amplitudes were larger during personalized strong than weak CST states and during personalized strong than random CST states (p < 0.03 for both) but did not differ between personalized weak and random CST states (p = 0.257). See Table 1 for percentage differences in MEP amplitudes elicited between states.

**Figure 3.**
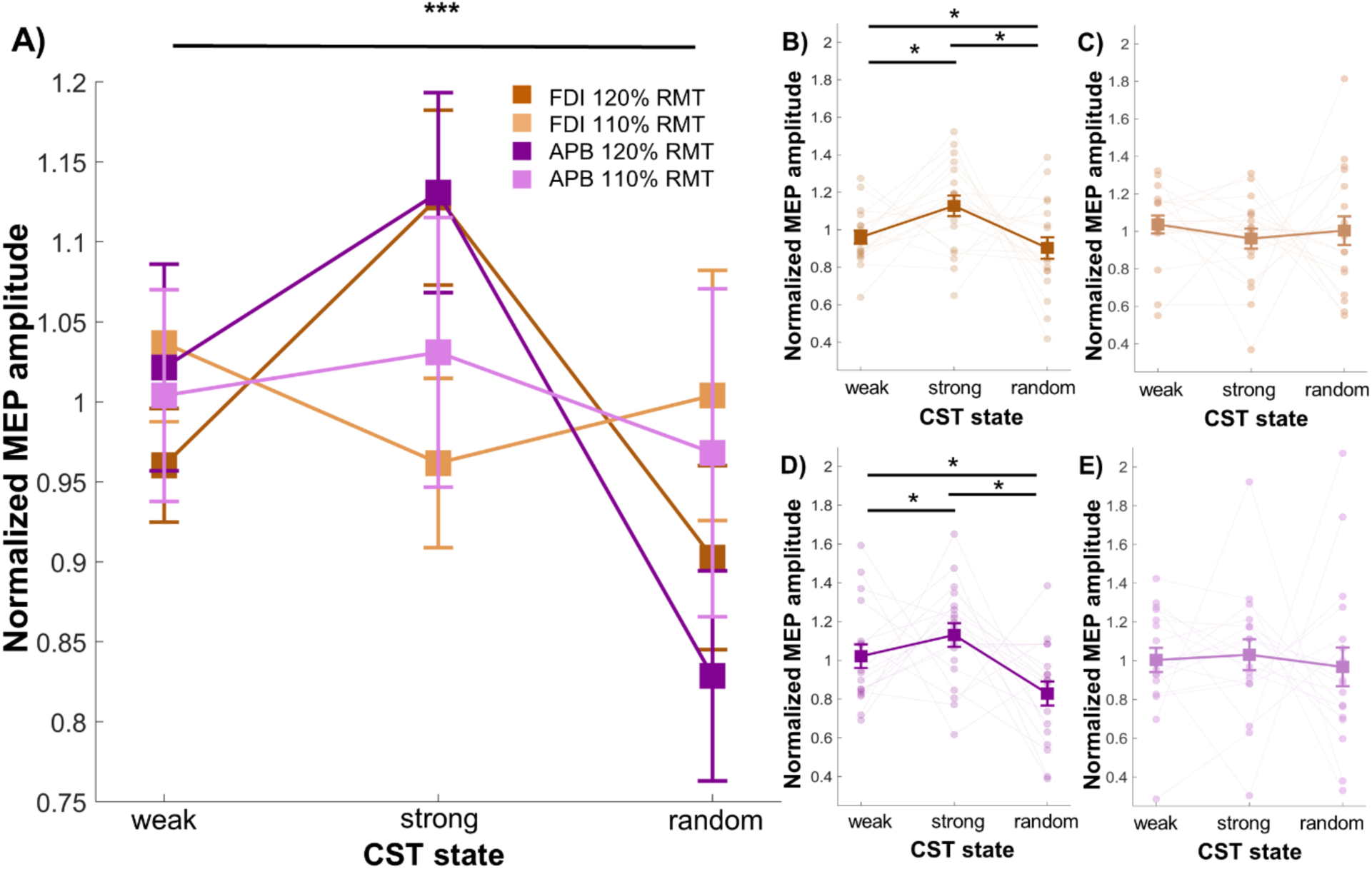
MEP amplitudes elicited during personalized strong, weak, and random CST states in real-time. (A) MEP amplitudes recorded from the L. FDI and L. APB muscles during real-time, classifier-predicted personalized strong, weak, and random CST states at 120% and 110% RMT. MEP amplitudes recorded from (B) L. FDI at 120% RMT, (C) L. FDI at 110% RMT, (D) L. APB at 120% RMT, (E) L. APB at 110% RMT. Triple asterisks reflect significant STATE x INTENSITY interaction. Single asterisks reflect significant pairwise post hoc comparisons for the STATE x INTENSITY interaction. Squares denote group averages, circles denote data from individual participants, and error bars reflect SEM.

**Table 1.**
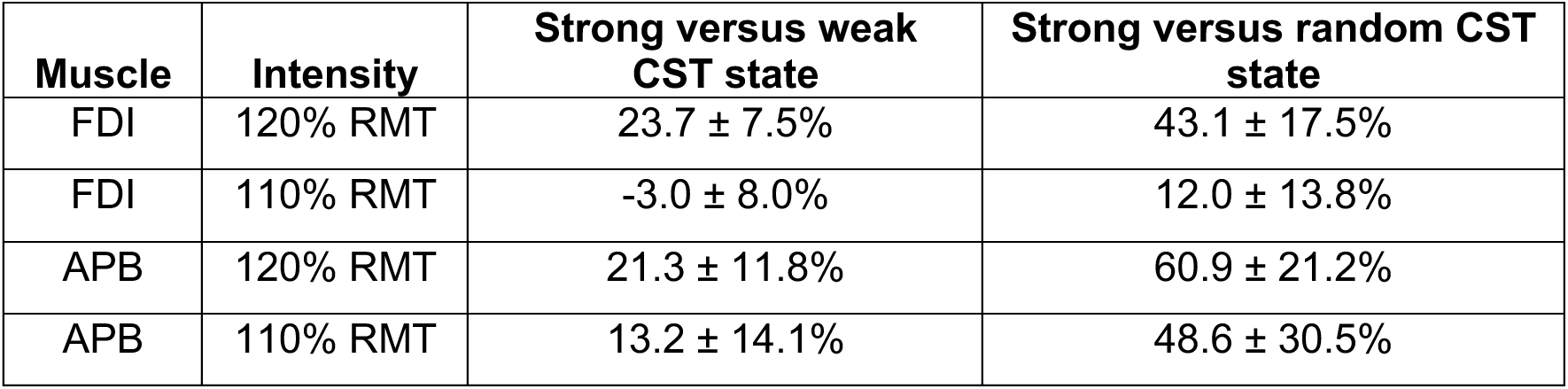
Percentage difference in MEP amplitudes elicited during personalized strong versus weak CST states and personalized strong versus random CST states for each muscle and stimulation intensity.

| Muscle | Intensity | Strong versus weak CST state | Strong versus random CST state |
| --- | --- | --- | --- |
| FDI | 120% RMT | $23.7 \pm 7.5\%$ | $43.1 \pm 17.5\%$ |
| FDI | 110% RMT | $-3.0 \pm 8.0\%$ | $12.0 \pm 13.8\%$ |
| APB | 120% RMT | $21.3 \pm 11.8\%$ | $60.9 \pm 21.2\%$ |
| APB | 110% RMT | $13.2 \pm 14.1\%$ | $48.6 \pm 30.5\%$ |

### MEP variability

We next examined MEP amplitude variability during personalized strong, weak, and random CST states by comparing coefficients of variation calculated from trial-by-trial MEP amplitudes across CST states at both intensities (120% and 110% RMT) and both muscles (L. FDI and L. APB, Figure 4). Linear-mixed effects models identified a significant main effect of STATE (F = 3.36, p = 0.037), INTENSITY (F = 87.6, p < 0.001), and MUSCLE (F = 52.03, p < 0.001). Post hoc pairwise comparisons showed that MEP amplitude variability was significantly lower during personalized strong than weak CST states (p = 0.04). However, MEP amplitude variability did not differ between personalized weak and random CST states or between personalized strong and random CST states (p > 0.07 for both). MEP amplitudes elicited at 120% RMT were less variable than those elicited at 110% RMT (p < 0.001) and MEP amplitudes recorded from L. FDI were less variable than those recorded from L. APB (p < 0.001).

**Figure 4.**
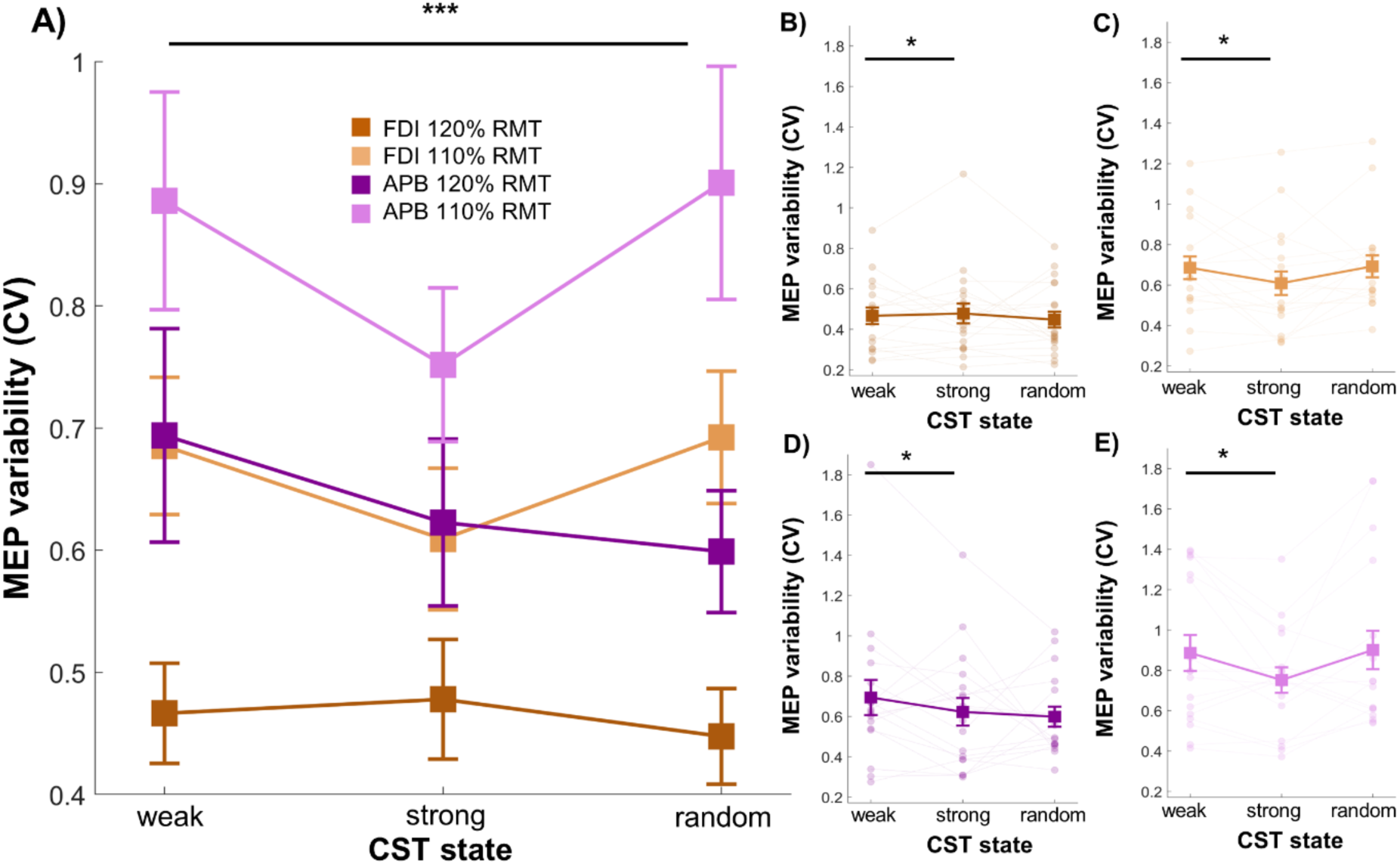
Variability of MEP amplitudes elicited during personalized strong, weak, and random CST states in real-time. (A) Coefficients of variation calculated from trial-by-trial MEP amplitudes recorded from the L. FDI and L. APB muscles during real-time, classifier-predicted personalized strong, weak, and random CST states at 120% and 110% RMT. Coefficients of variation calculated from trial-by-trial MEP amplitudes recorded from (B) L. FDI at 120% RMT, (C) L. FDI at 110% RMT, (D) L. APB at 120% RMT, and (E) L. APB at 110% RMT. Triple asterisks reflect the significant main effects of STATE, MUSCLE and INTENSITY, and single asterisks reflect significant pairwise post hoc comparisons for the main effect of STATE. Squares denote group averages, circles denote data from individual participants, and error bars denote SEM.

### Temporal characteristics of CST states

We also evaluated the temporal characteristics of personalized strong and weak CST states by applying each participant’s classifier to their resting EEG recording. When examining the percentage of time during which either strong, weak, or under-confident CST states were detected by personalized classifiers (Figure 5A), a Kruskal-Wallis test revealed a significant effect of STATE (p = 0.002). Post hoc pairwise comparisons showed that the proportion of time spent per state did not differ between strong and weak CST states (p = 0.125) but was significantly lower for under-confident than weak or strong CST states (p < 0.027 for both). Overall, personalized strong CST states were present 33.6 ± 5.9% of the time, personalized weak CST states were present 48.4 ± 6.5% of the time, and under-confident CST states were present 17.8 ± 2.6% of the time.

**Figure 5.**
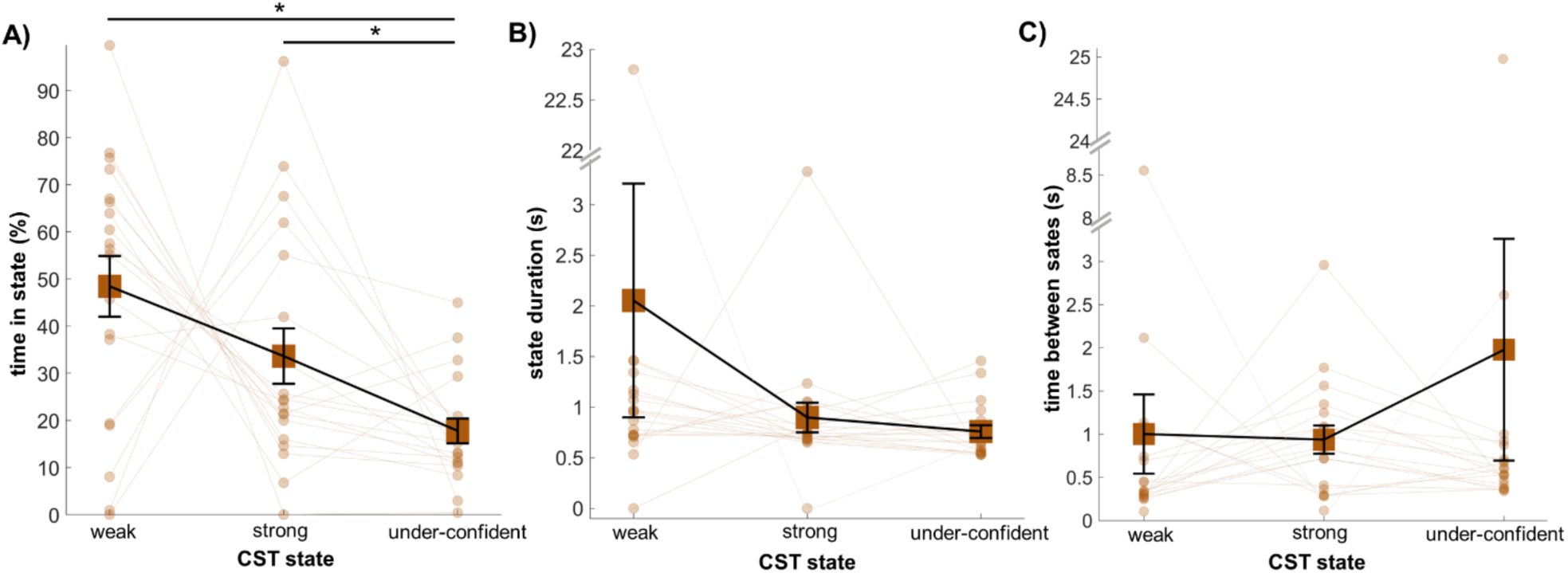
Temporal characteristics of personalized CST states present during resting EEG recordings. (A) Percentage of time spent in each CST state. (B) Average duration of each CST state. C) Time between consecutive CST states. Squares denote group averages, circles denote data from individual participants, and error bars denote SEM. Asterisks indicate significant post hoc pairwise comparisons between CST states.

When evaluating the mean duration of strong, weak, or under-confident CST states detected by personalized classifiers (Figure 5B), a Kruskal-Wallis test did not show any significant effect of STATE. Personalized strong CST states lasted on average 0.89 ± 0.14 s, personalized weak CST states lasted on average 2.05 ± 1.15 s, and under-confident CST states lasted on average 0.75 ± 0.06 s.

When examining the mean time between consecutive strong, weak, or under-confident CST states detected by personalized classifiers (Figure 5C), a Kruskal-Wallis test did not reveal a significant effect of STATE. On average, 0.93 ± 0.16 s elapsed between consecutive strong CST states, 1.0 ± 0.45 s elapsed between consecutive weak CST states, and 1.97 ± 1.28 s elapsed between consecutive under-confident CST states.

### Personalized versus non-personalized classifier performance

Personalized classifier F1 values were on average 0.68 ± 0.01, while non-personalized classifier F1 values were on average 0.69 ± 0.01. F1 values did not differ across classifier types (p = 0.33; see Supplementary Figure 1).

### Relationships between personalized classifier performance and brain state-dependent variation in MEP amplitudes

Overall, L. FDI MEP amplitudes elicited at 120% RMT during personalized strong CST states were 23.7 ± 7.5% and 43.1 ± 17.5% larger than those elicited during personalized weak and random CST states, respectively (see Table 1). The percentage difference in MEP amplitude between CST states did not correlate with personalized F1 values (percentage difference between strong and weak CST states versus F1 values: R = 0.14, p = 0.56; percentage difference between strong and random CST states versus F1 values: R = 0.12, p = 0.63; see Supplementary Figure 2).

### Spectro-spatial characteristics of personalized CST states

We characterized the spectro-spatial characteristics of pre-stimulus EEG activity present during real-time classifier-predicted personalized strong versus weak CST states. At the individual level, each participant exhibited qualitatively unique differences in pre-stimulus EEG power between personalized strong and weak CST states across the scalp (see Supplementary Figure 3). For example, participant #13 showed higher right parieto-occipital alpha power during strong than weak CST states, while participant #14 exhibited higher right frontal beta power and lower whole-scalp theta power during strong than weak CST states. At the group-level, right centro-parietal alpha power and whole-scalp theta power were generally higher during personalized strong than weak CST states, and TP9 and TP10 showed stronger theta, alpha, and beta power during strong versus weak CST states.

**Figure 6.**
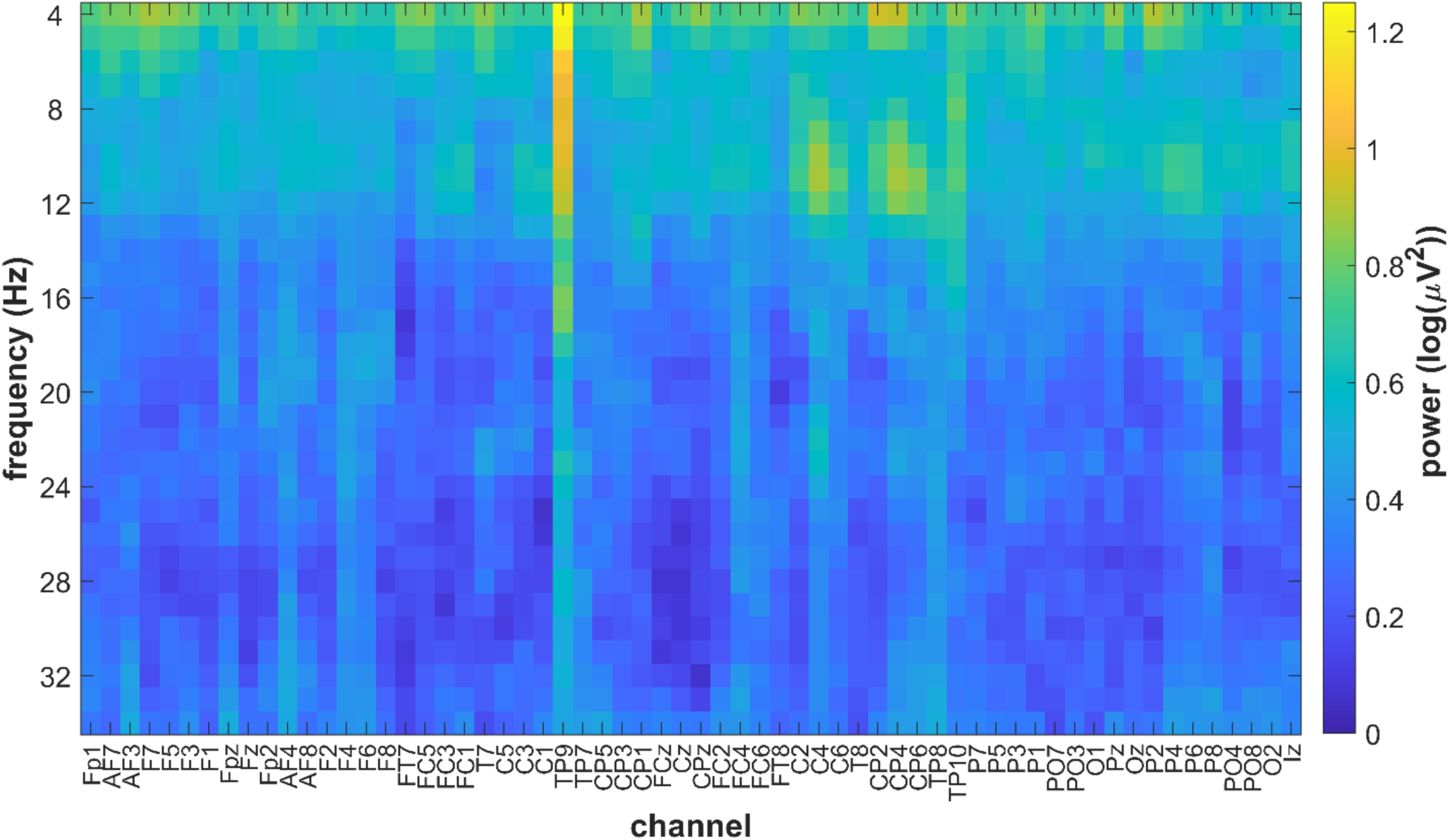
Group-level differences in pre-stimulus spectro-spatial EEG patterns corresponding to personalized strong and weak CST states identified in real-time. Group average differences in natural log-transformed power between EEG activity during personalized strong and weak CST states. See Supplementary Figure 3 for individual participant data. Note that dark blue values reflect 0, which indicates no difference in power between states.

## Discussion

In this study, we developed a first-of-its-kind machine learning-driven real-time EEG-TMS system that delivers TMS during personalized brain activity patterns reflecting strong and weak CST activation in healthy humans at rest. We report that this system accurately targets personalized strong and weak CST states, such that MEPs elicited during personalized strong CST states were significantly larger than those elicited during personalized weak and random CST states. Although this pattern of results was present for both L. FDI and L. APB muscles, it was only evident when evaluating the same stimulation intensity used to train personalized classifiers (i.e., 120% RMT). Additionally, personalized strong and weak CST states were present ∼35% and ∼50% of the time, respectively, and typically lasted for ∼1-2 seconds. Group-level spectro-spatial differences in pre-stimulus EEG activity showed that whole-scalp theta power and right centro-parietal alpha power were higher during personalized strong than weak CST states. Overall, our results demonstrate the feasibility and efficacy of real-time personalized brain state-dependent TMS targeting the human CST and are a key step towards future interventional studies using this novel decoding-based brain stimulation technique. Recent studies have shown that MEP amplitudes are ∼10-20% larger during optimal than nonoptimal sensorimotor rhythm phases (Bergmann et al., 2019; Ozdemir et al., 2022; Suresh & Hussain, 2023; Zrenner et al., 2018), with some studies reporting no difference (Madsen et al., 2019b). In the current study, we trained personalized classifiers to discriminate between EEG patterns during which TMS elicited large and small MEPs (i.e., personalized strong and weak CST states) using single-pulse TMS-EEG-EMG datasets acquired from L. FDI at 120% RMT. MEPs elicited in real-time at this same muscle and intensity were ∼24% and ∼43% larger than those elicited in real-time during corresponding weak and random CST states, respectively. The magnitude of state-dependent MEP amplitude variation observed here exceeds that reported in previous phase-dependent single-pulse TMS studies. MEPs elicited from the L. APB muscle at 120% RMT showed a similar pattern of results (see Figure 4), indicating that classifiers trained to identify personalized strong and weak CST states generalize across intrinsic hand muscles. Given that the two muscles evaluated here are both functionally related (i.e., involved in grasping behaviors) and topographically adjacent within the sensorimotor cortex, the personalized CST states targeted here either capture dynamic fluctuations in excitability of functionally-coupled cortical muscle representations, spatially-coupled cortical muscle representations, or both. In contrast, classifiers trained using TMS-EEG-EMG datasets acquired at 120% RMT did not reliably elicit larger MEPs in real-time during personalized strong versus weak or random CST states at 110% RMT. That is, personalized classifiers did not generalize across stimulation intensities. This lack of generalization may be due to differences in the motor cortical interneuronal circuits activated by TMS at 120% and 110% RMT. Given that higher intensity TMS elicits a greater number of indirect waves as well as direct waves (Lazzaro et al., 2014), the personalized EEG patterns identified from our training dataset may be specific to the precise combination of descending corticospinal volleys elicited at 120% RMT. Future studies could improve the flexibility and generalizability of machine learning-driven real-time EEG-TMS by training personalized classifiers on TMS-EEG-EMG datasets acquired from multiple muscles at multiple intensities.

To date, studies examining brain state-dependency of CST activation have either not reported trial-by-trial variation in MEP amplitudes across different brain states (Bergmann et al., 2019; Hussain et al., 2019; Thies et al., 2018; Wischnewski et al., 2022; Zrenner et al., 2018) or identified no differences (Ozdemir et al., 2022). Here, we show brain state-dependency of MEP amplitude variability for the first time, reporting that trial-by-trial variation in MEP amplitudes is significantly lower during personalized strong than weak CST states. In addition to more strongly activating the CST, these findings suggest that TMS also more consistently activates the CST when delivered during personalized strong CST states, which may benefit effect sizes of future TMS interventions targeting these states. Surprisingly, however, MEP amplitude variability did not differ between strong and random CST states, nor did it differ between weak and random CST states. This may be because we evaluated MEP amplitudes during random CST states using conventional brain state-independent TMS, rather than a mixture of high and low CST states. Consistent with reports that MEP amplitudes are less variable at higher stimulation intensities (Darling et al., 2006; Schaworonkow et al., 2019), we observed that MEPs were less variable at 120% than 110% RMT. Finally, MEPs recorded from L. APB were more variable than those recorded from L. FDI, likely because the scalp TMS site was optimized based on L. FDI rather than L. APB responses.

The defining feature of brain state-dependent TMS interventions is that individual TMS pulses are only delivered when the desired brain activity pattern is detected in real-time. As a result, the brain states targeted by such interventions must occur frequently enough that the desired number of TMS pulses can be delivered within a feasible timeframe. When delivering real-time single-pulse TMS during personalized strong and weak CST states in the current study, inter-stimulus intervals ranged on average between ∼7-10 seconds. In addition to the frequency with which personalized CST states occurred, these inter-stimulus intervals are directly influenced by the minimum allowable inter-stimulus interval (here, 3 seconds) and the number of consecutive CST states our real-time EEG analysis system required before TMS delivery (here, 10 consecutive states). To better quantify the temporal characteristics of personalized strong and weak CST states without these methodological constraints, we applied each participant’s personalized classifier to their resting EEG data using a sliding window approach. This analysis revealed that the temporal characteristics of personalized strong and weak CST states did not differ. On average, personalized strong and weak CST states were present 35-50% of the time and lasted for ∼1-2 seconds; ∼1 second elapsed between consecutive similar CST states. Thus, personalized strong and weak CST states appear to be sufficiently frequent and adequately prolonged to be targeted with repeated TMS pulses during an intervention. To finely tune inter-stimulus intervals during TMS interventions, future studies can modify the number of consecutive CST states that the real-time EEG analysis algorithm must detect before triggering TMS pulses.

A major advantage of our approach is that it requires no prior knowledge regarding which EEG activity patterns reflect strong versus weak CST activation in each participant. To characterize the spectro-spatial EEG patterns present during personalized strong and weak CST states, we performed participant-specific contrasts between EEG power spectral activity recorded during real-time targeting of strong and weak states. Consistent with our personalized approach, these contrasts identified unique whole-scalp EEG activity patterns that discriminated between personalized strong and weak CST states in individual participants (see Supplementary Figure 3 for details). Given that whole-scalp EEG signals likely capture the influence of long-range inter- and intra-hemispheric projections to motor cortical interneurons and CST neurons, whole-scalp EEG may be particularly useful for identifying personalized CST states. For example, previous work suggests that long-range projections can modulate corticospinal output (Bestmann & Krakauer, 2015), including those originating from the supplementary motor area (Arai et al., 2012), premotor cortex (Münchau et al., 2002), dorsolateral prefrontal cortex (Hasan et al., 2013), and cerebellum and basal ganglia via thalamic nuclei (Sommer, 2003). Group-level spectro-spatial characteristics revealed that right centro-parietal alpha power and whole-scalp theta power were generally higher during personalized strong than weak CST states. The centro-parietal alpha activity identified in the current study is broadly consistent with recent reports that corticospinal output rhythmically fluctuates at alpha frequencies (Metsomaa et al., 2021) and relates to alpha activity near the stimulated cortex and parieto-occipital regions (Ermolova et al., 2024). The consistency in these group-level findings across studies suggest that the use of personalized classifiers to identify strong and weak CST states may not be truly necessary, at least in healthy adults. To explore this possibility, we built a single non-personalized classifier that could discriminate between strong and weak CST states using TMS-EEG-EMG training datasets acquired from all participants. Surprisingly, the performance of personalized and non-personalized classifiers did not differ, suggesting that personalization may not be essential for brain state-dependent TMS in healthy adults. However, it is important to note that the non-personalized classifier was trained using substantially more data than personalized classifiers, which likely improved its performance. Further, this non-personalized classifier’s ability to accurately identify CST states in real-time remains untested. Finally, a key goal of brain state-dependent TMS is to improve the therapeutic efficacy of poststroke TMS interventions. Given the heterogeneity of lesion characteristics (Chen et al., 2000; Luft et al., 2004; Shelton & Reding, 2001), recovery-related adaptive plasticity (Grefkes & Ward, 2014; Stinear et al., 2007), and alterations in sensorimotor oscillatory dynamics after stroke (Hussain et al., 2020; Johnston et al., 2023; Lopez-Larraz et al., 2017), personalized classifiers may be essential for accurate poststroke brain state-dependent TMS.

Limitations to this study also exist. First, therapeutic TMS interventions are often delivered over multiple days, but the between-day generalizability of personalized classifiers developed here has yet to be tested. Our method requires a large, participant-specific TMS-EEG-EMG training dataset to build each participant’s unique classifier and acquiring a new training dataset on each treatment day is likely not clinically feasible. However, between-participant generalization of machine learning classifiers used for brain-computer interfaces can be improved by using advanced statistical matching procedures that do not require any calibration for each participant (Kumar et al., 2024). Similar procedures could be applied to improve the between-day generalizability of personalized classifiers developed here. Second, some participants showed poor state targeting accuracy for weak CST states, suggesting that weak states may be less reliable than strong CST states. Although most therapeutic applications of the machine learning-driven TMS approach developed here are likely to focus on increasing CST transmission by targeting strong CST states, the stability and persistence of weak CST states requires further investigation. Finally, the performance of the personalized classifiers developed here is lower than reported in conventional brain-computer interface paradigms but is consistent with multiple recent studies that used machine learning to identify EEG patterns reflecting strong and weak CST states, both from our group (Hussain et al., 2022) and others (Ermolova et al., 2024; Metsomaa et al., 2021). The overall higher performance of brain-computer interface classifiers compared to our approach likely relates to the volitional modulation of EEG signals in such contexts, typically via motor imagery (Perdikis & Millan, 2020; Tonin et al., 2022). Further, we observed that classifier performance did not correlate with the difference in CST activation between personalized strong and weak CST states. This lack of relationship may be caused by variation in spinal motoneuron depolarization that is not represented within EEG signals. Regardless, brain state-dependent TMS interventions are most effective when delivered during EEG activity patterns associated with large MEPs (Baur et al., 2020; Zrenner et al., 2018), indicating that MEP amplitude differences between targeted CST states are likely more important for inducing strong neuroplastic effects than classifier performance *per se*.

In conclusion, here we demonstrate for the first time that personalized brain activity patterns reflecting strong and weak CST activation can be accurately captured in real-time using machine learning-driven whole scalp EEG-triggered TMS in healthy adults. Specifically, we report that CST activation was greater during personalized strong than weak and random CST states and was also more consistent during personalized strong than weak CST states. Personalized strong and weak CST states lasted for ∼1-2 seconds at a time and ∼1 second elapsed between similar consecutive states, suggesting that personalized CST states could be repeatedly targeted with TMS during future interventional applications. Individual participants also exhibited unique spectro-spatial EEG patterns that differed between strong and weak CST states; these patterns are likely to be even more heterogeneous poststroke. Overall, our findings represent a key step towards using personalized brain state-dependent TMS techniques to characterize and promote poststroke CST function.

## Acknowledgments

The research reported in this publication was supported by the National Institute of Neurological Disorders and Stroke of the National Institutes of Health under Award Number R21NS133605. SJH was supported by K12HD0903427, R21NS133605, and R03HD114118. UUK and JdRM were supported by R21NS133605. The content is solely the responsibility of the authors and does not necessarily represent the official views of the National Institutes of Health.

## Data availability

Raw data and analysis code will be shared via an open-access repository prior to acceptance for publication.

## References

Arai, N., Lu, M. K., Ugawa, Y., & Ziemann, U. (2012). Effective connectivity between human supplementary motor area and primary motor cortex: A paired-coil TMS study. Experimental Brain Research, 220(1), 79–87. 10.1007/S00221-012-3117-5/TABLES/1

Awiszus, F. (2011). Fast estimation of transcranial magnetic stimulation motor threshold: Is it safe? Brain Stimulation, 4(1), 50–57. 10.1016/J.BRS.2010.06.002

Baur, D., Galevska, D., Hussain, S., Cohen, L. G., Ziemann, U., & Zrenner, C. (2020). Induction of LTD-like corticospinal plasticity by low-frequency rTMS depends on pre-stimulus phase of sensorimotor μ-rhythm. Brain Stimulation, 13(6), 1580–1587. 10.1016/J.BRS.2020.09.005

Benjamini, Y., & Hochberg, Y. (1995). Controlling the False Discovery Rate: A Practical and Powerful Approach to Multiple Testing. Journal of the Royal Statistical Society: Series B (Methodological), 57(1), 289–300. 10.1111/J.2517-6161.1995.TB02031.X

Berger, B., Minarik, T., Liuzzi, G., Hummel, F. C., & Sauseng, P. (2014). EEG Oscillatory Phase-Dependent Markers of Corticospinal Excitability in the Resting Brain. BioMed Research International, 2014(1), 936096. 10.1155/2014/936096

Bergmann, T. O., Lieb, A., Zrenner, C., & Ziemann, U. (2019). Pulsed facilitation of corticospinal excitability by the sensorimotor μ-alpha rhythm. Journal of Neuroscience, 39(50), 10034–10043. 10.1523/JNEUROSCI.1730-19.2019

Bestmann, S., & Krakauer, J. W. (2015). The uses and interpretations of the motor-evoked potential for understanding behaviour. Experimental Brain Research, 233(3), 679–689. 10.1007/S00221-014-4183-7/METRICS

Blankertz, B., Tomioka, R., Lemm, S., Kawanabe, M., & Müller, K. R. (2008). Optimizing spatial filters for robust EEG single-trial analysis. IEEE Signal Processing Magazine, 25(1), 41–56. 10.1109/MSP.2008.4408441

Bunday, K. L., & Perez, M. A. (2012). Motor recovery after spinal cord injury enhanced by strengthening corticospinal synaptic transmission. Current Biology, 22(24), 2355–2361. 10.1016/j.cub.2012.10.046

Chen, C. L., Tang, F. T., Chen, H. C., Chung, C. Y., & Wong, M. K. (2000). Brain lesion size and location: Effects on motor recovery and functional outcome in stroke patients. Archives of Physical Medicine and Rehabilitation, 81(4), 447–452. 10.1053/MR.2000.3837

Chen, R., Tam, A., Bütefisch, C., Corwell, B., Ziemann, U., Rothwell, J. C., & Cohen, L. G. (1998). Intracortical inhibition and facilitation in different representations of the human motor cortex. Journal of Neurophysiology, 80(6), 2870–2881. 10.1152/JN.1998.80.6.2870

Darling, W. G., Wolf, S. L., & Butler, A. J. (2006). Variability of motor potentials evoked by transcranial magnetic stimulation depends on muscle activation. Experimental Brain Research, 174(2), 376–385. 10.1007/S00221-006-0468-9/TABLES/1

Delvaux, V., Alagona, G., Gérard, P., De Pasqua, V., Pennisi, G., & De Noordhout, A. M. (2003). Post-stroke reorganization of hand motor area: a 1-year prospective follow-up with focal transcranial magnetic stimulation. Clinical Neurophysiology, 114(7), 1217–1225. 10.1016/S1388-2457(03)00070-1

Di Lazzaro, V., Pilato, F., Dileone, M., Profice, P., Capone, F., Ranieri, F., Musumeci, G., Cianfoni, A., Pasqualetti, P., & Tonali, P. A. (2008). Modulating cortical excitability in acute stroke: A repetitive TMS study. Clinical Neurophysiology, 119(3), 715–723. 10.1016/J.CLINPH.2007.11.049

Di Lazzaro, V., & Ziemann, U. (2013). The contribution of transcranial magnetic stimulation in the functional evaluation of microcircuits in human motor cortex. In Frontiers in Neural Circuits (Issue JAN). 10.3389/fncir.2013.00018

Du, J., Tian, L., Liu, W., Hu, J., Xu, G., Ma, M., Fan, X., Ye, R., Jiang, Y., Yin, Q., Zhu, W., Xiong, Y., Yang, F., & Liu, X. (2016). Effects of repetitive transcranial magnetic stimulation on motor recovery and motor cortex excitability in patients with stroke: a randomized controlled trial. European Journal of Neurology, 23(11), 1666–1672. 10.1111/ENE.13105

Ermolova, M., Metsomaa, J., Belardinelli, P., Zrenner, C., & Ziemann, U. (2024). Blindly separated spontaneous network-level oscillations predict corticospinal excitability. Journal of Neural Engineering, 21(3), 036041. 10.1088/1741-2552/AD5404

Freedberg, M. V., Reeves, J. A., Fioriti, C. M., Murillo, J., & Wassermann, E. M. (2022). Reproducing the effect of hippocampal network-targeted transcranial magnetic stimulation on episodic memory. Behavioural Brain Research, 419, 113707. 10.1016/J.BBR.2021.113707

George, M. S., Lisanby, S. H., Avery, D., McDonald, W. M., Durkalski, V., Pavlicova, M., Anderson, B., Nahas, Z., Bulow, P., Zarkowski, P., Holtzheimer, P. E., Schwartz, T., & Sackeim, H. A. (2010). Daily Left Prefrontal Transcranial Magnetic Stimulation Therapy for Major Depressive Disorder: A Sham-Controlled Randomized Trial. Archives of General Psychiatry, 67(5), 507–516. 10.1001/ARCHGENPSYCHIATRY.2010.46

George, M. S., Wassermann, E. M., Williams, W. A., Callahan, A., Ketter, T. A., Basser, P., Hallett, M., & Post, R. M. (1995). Daily repetitive transcranial magnetic stimulation (rTMS) improves mood in depression. Neuroreport, 6(14), 1853–1856. 10.1097/00001756-199510020-00008

Grefkes, C., & Ward, N. S. (2014). Cortical reorganization after stroke: how much and how functional? *The Neuroscientist : A Review Journal Bringing Neurobiology*, Neurology and Psychiatry, 20(1), 56–70. 10.1177/1073858413491147

Hamada, M., Murase, N., Hasan, A., Balaratnam, M., & Rothwell, J. C. (2013). The Role of Interneuron Networks in Driving Human Motor Cortical Plasticity. Cerebral Cortex, 23(7), 1593–1605. 10.1093/CERCOR/BHS147

Hasan, A., Galea, J. M., Casula, E. P., Falkai, P., Bestmann, S., & Rothwell, J. C. (2013). Muscle and Timing-specific Functional Connectivity between the Dorsolateral Prefrontal Cortex and the Primary Motor Cortex. Journal of Cognitive Neuroscience, 25(4), 558–570. 10.1162/JOCN_A_00338

Hoogendam, J. M., Ramakers, G. M. J., & Di Lazzaro, V. (2010). Physiology of repetitive transcranial magnetic stimulation of the human brain. Brain Stimulation, 3(2), 95–118. 10.1016/J.BRS.2009.10.005

Huang, Y. Z., Edwards, M. J., Rounis, E., Bhatia, K. P., & Rothwell, J. C. (2005). Theta burst stimulation of the human motor cortex. Neuron, 45(2), 201–206. 10.1016/j.neuron.2004.12.033

Hussain, S. J., Claudino, L., Bönstrup, M., Norato, G., Cruciani, G., Thompson, R., Zrenner, C., Ziemann, U., Buch, E., & Cohen, L. G. (2019a). Sensorimotor Oscillatory Phase–Power Interaction Gates Resting Human Corticospinal Output. Cerebral Cortex, 29(9), 3766– 3777. 10.1093/CERCOR/BHY255

Hussain, S. J., Claudino, L., Bönstrup, M., Norato, G., Cruciani, G., Thompson, R., Zrenner, C., Ziemann, U., Buch, E., & Cohen, L. G. (2019b). Sensorimotor Oscillatory Phase–Power Interaction Gates Resting Human Corticospinal Output. Cerebral Cortex, 29(9), 3766– 3777. 10.1093/CERCOR/BHY255

Hussain, S. J., Hayward, W., Fourcand, F., Zrenner, C., Ziemann, U., Buch, E. R., Hayward, M. K., & Cohen, L. G. (2020). Phase-dependent transcranial magnetic stimulation of the lesioned hemisphere is accurate after stroke. Brain Stimulation, 13(5), 1354–1357. 10.1016/j.brs.2020.07.005

Hussain, S. J., Quentin, R., Lyon, B., & Lyon, F. (2022). Decoding personalized motor cortical excitability states from human electroencephalography. Scientific Reports *2022 12:1*, 12(1), 1–12. 10.1038/s41598-022-10239-3

Jo, H. J., & Perez, M. A. (2020). Corticospinal-motor neuronal plasticity promotes exercise-mediated recovery in humans with spinal cord injury. Brain, 143(5), 1368–1382. 10.1093/BRAIN/AWAA052

Johnston, P. R., McIntosh, A. R., & Meltzer, J. A. (2023). Spectral slowing in chronic stroke reflects abnormalities in both periodic and aperiodic neural dynamics. NeuroImage. Clinical, 37. 10.1016/J.NICL.2022.103277

Jones, T. A. (2017). Motor compensation and its effects on neural reorganization after stroke. Nature Reviews Neuroscience *2017 18:5*, 18(5), 267–280. 10.1038/nrn.2017.26

Jung, N. H., Delvendahl, I., Kuhnke, N. G., Hauschke, D., Stolle, S., & Mall, V. (2010). Navigated transcranial magnetic stimulation does not decrease the variability of motor-evoked potentials. Brain Stimulation, 3(2), 87–94. 10.1016/J.BRS.2009.10.003

Kiers, L., Cros, D., Chiappa, K. H., & Fang, J. (1993). Variability of motor potentials evoked by transcranial magnetic stimulation. Electroencephalography and Clinical Neurophysiology/Evoked Potentials Section, 89(6), 415–423. 10.1016/0168-5597(93)90115-6

Kothe, C., Shirazi, S. Y., Stenner, T., Medine, D., Boulay, C., Grivich, M. I., Mullen, T., Delorme, A., & Makeig, S. (2024). The Lab Streaming Layer for Synchronized Multimodal Recording. BioRxiv, 2024.02.13.580071. 10.1101/2024.02.13.580071

Kumar, S., Alawieh, H., Racz, F. S., Fakhreddine, R., & del Millán, J. R. (2024). Transfer learning promotes acquisition of individual BCI skills. PNAS Nexus, 3(2). 10.1093/PNASNEXUS/PGAE076

Lazzaro, V. Di, Rothwell, J. C., Lazzaro, V. Di, & Rothwell, J. C. (2014). Corticospinal activity evoked and modulated by non-invasive stimulation of the intact human motor cortex. The Journal of Physiology, 592(19), 4115–4128. 10.1113/JPHYSIOL.2014.274316

Lemon, R. N. (2008). Descending pathways in motor control. Annual Review of Neuroscience, 31(Volume 31, 2008), 195–218. 10.1146/ANNUREV.NEURO.31.060407.125547/CITE/REFWORKS

López-Alonso, V., Cheeran, B., Río-Rodríguez, D., & Fernández-Del-Olmo, M. (2014). Inter-individual variability in response to non-invasive brain stimulation paradigms. Brain Stimulation, 7(3), 372–380. 10.1016/J.BRS.2014.02.004

Lopez-Larraz, E., Ray, A. M., Figueiredo, T. C., Bibian, C., Birbaumer, N., & Ramos-Murguialday, A. (2017). Stroke lesion location influences the decoding of movement intention from EEG. Proceedings of the Annual International Conference of the IEEE Engineering in Medicine and Biology Society, EMBS, 3065–3068. 10.1109/EMBC.2017.8037504

Lotze, M., Beutling, W., Loibl, M., Domin, M., Platz, T., Schminke, U., & Byblow, W. D. (2012). Contralesional motor cortex activation depends on ipsilesional corticospinal tract integrity in well-recovered subcortical stroke patients. Neurorehabilitation and Neural Repair, 26(6). 10.1177/1545968311427706

Luber, B., & Lisanby, S. H. (2014). Enhancement of human cognitive performance using transcranial magnetic stimulation (TMS). NeuroImage, 85, 961–970. 10.1016/J.NEUROIMAGE.2013.06.007

Luft, A. R., Waller, S., Forrester, L., Smith, G. V., Whitall, J., Macko, R. F., Schulz, J. B., & Hanley, D. F. (2004). Lesion location alters brain activation in chronically impaired stroke survivors. NeuroImage, 21(3), 924–935. 10.1016/J.NEUROIMAGE.2003.10.026

Madsen, K. H., Karabanov, A. N., Krohne, L. G., Safeldt, M. G., Tomasevic, L., & Siebner, H. R. (2019a). No trace of phase: Corticomotor excitability is not tuned by phase of pericentral mu-rhythm. Brain Stimulation, 12(5), 1261–1270. 10.1016/J.BRS.2019.05.005

Madsen, K. H., Karabanov, A. N., Krohne, L. G., Safeldt, M. G., Tomasevic, L., & Siebner, H. R. (2019b). No trace of phase: Corticomotor excitability is not tuned by phase of pericentral mu-rhythm. Brain Stimulation, 12(5), 1261–1270. 10.1016/J.BRS.2019.05.005

Mantovani, A., Lisanby, S. H., Pieraccini, F., Ulivelli, M., Castrogiovanni, P., & Rossi, S. (2006). Repetitive transcranial magnetic stimulation (rTMS) in the treatment of obsessive– compulsive disorder (OCD) and Tourette’s syndrome (TS). International Journal of Neuropsychopharmacology, 9(1), 95–100. 10.1017/S1461145705005729

Metsomaa, J., Belardinelli, P., Ermolova, M., Ziemann, U., & Zrenner, C. (2021). Causal decoding of individual cortical excitability states. NeuroImage, 245, 118652. 10.1016/J.NEUROIMAGE.2021.118652

Mills, K. R., Boniface, S. J., & Schubert, M. (1992). Magnetic brain stimulation with a double coil: the importance of coil orientation. Electroencephalography and Clinical Neurophysiology/Evoked Potentials Section, 85(1), 17–21. 10.1016/0168-5597(92)90096-T

Münchau, A., Bloem, B. R., Irlbacher, K., Trimble, M. R., & Rothwell, J. C. (2002). Functional Connectivity of Human Premotor and Motor Cortex Explored with Repetitive Transcranial Magnetic Stimulation. Journal of Neuroscience, 22(2), 554–561. 10.1523/JNEUROSCI.22-02-00554.2002

Oostenveld, R., Fries, P., Maris, E., & Schoffelen, J. M. (2011). FieldTrip: Open Source Software for Advanced Analysis of MEG, EEG, and Invasive Electrophysiological Data. Computational Intelligence and Neuroscience, 2011(1), 156869. 10.1155/2011/156869

Ozdemir, R. A., Kirkman, S., Magnuson, J. R., Fried, P. J., Pascual-Leone, A., & Shafi, M. M. (2022). Phase matters when there is power: Phasic modulation of corticospinal excitability occurs at high amplitude sensorimotor mu-oscillations. Neuroimage: Reports, 2(4), 100132. 10.1016/J.YNIRP.2022.100132

Park, W., Kwon, G. H., Kim, Y. H., Lee, J. H., & Kim, L. (2016). EEG response varies with lesion location in patients with chronic stroke. Journal of NeuroEngineering and Rehabilitation, 13(1), 1–10. 10.1186/S12984-016-0120-2/FIGURES/4

Pascual-Leone, A., Dang, N., Cohen, L. G., Brasil-Neto, J. P., Cammarota, A., & Hallett, M. (1995). Modulation of Muscle Responses Evoked by Transcranial Magnetic Stimulation During the Acquisition of New Fine Motor Skills. JOURNALOFNEUROPHYSIOLOGY, 74(3).

Pellicciari, M. C., Miniussi, C., Ferrari, C., Koch, G., & Bortoletto, M. (2016). Ongoing cumulative effects of single TMS pulses on corticospinal excitability: An intra- and inter-block investigation. Clinical Neurophysiology, 127(1), 621–628. 10.1016/J.CLINPH.2015.03.002

Perdikis, S., & Millan, J. del R. (2020). Brain-Machine Interfaces: A Tale of Two Learners. IEEE Systems, Man, and Cybernetics Magazine, 6(3), 12–19. 10.1109/MSMC.2019.2958200

Schaworonkow, N., Triesch, J., Ziemann, U., & Zrenner, C. (2019). EEG-triggered TMS reveals stronger brain state-dependent modulation of motor evoked potentials at weaker stimulation intensities. Brain Stimulation, 12(1), 110–118. 10.1016/J.BRS.2018.09.009

Shelton, F. D. N. A. P., & Reding, M. J. (2001). Effect of lesion location on upper limb motor recovery after stroke. Stroke, 32(1), 107–112. 10.1161/01.STR.32.1.107/ASSET/4155D182-5B81-4877-A2CB-5564C8987EEB/ASSETS/GRAPHIC/HS0112242001.JPEG

Solé-Padullés, C., Bartrés-Faz, D., Junqué, C., Clemente, I. C., Molinuevo, J. L., Bargalló, N., Sánchez-Aldeguer, J., Bosch, B., Falcón, C., & Valls-Solé, J. (2006). Repetitive Transcranial Magnetic Stimulation Effects on Brain Function and Cognition among Elders with Memory Dysfunction. A Randomized Sham-Controlled Study. Cerebral Cortex, 16(10), 1487–1493. 10.1093/CERCOR/BHJ083

Sommer, M. A. (2003). The role of the thalamus in motor control. Current Opinion in Neurobiology, 13(6), 663–670. 10.1016/J.CONB.2003.10.014

Stinear, C. (2010). Prediction of recovery of motor function after stroke. The Lancet Neurology, 9(12), 1228–1232. 10.1016/S1474-4422(10)70247-7

Stinear, C. M., Barber, P. A., Smale, P. R., Coxon, J. P., Fleming, M. K., & Byblow, W. D. (2007). Functional potential in chronic stroke patients depends on corticospinal tract integrity. Brain, 130(1), 170–180. 10.1093/BRAIN/AWL333

Suresh, T., & Hussain, S. J. (2023). Re-evaluating the contribution of sensorimotor mu rhythm phase and power to human corticospinal output: A replication study. Brain Stimulation, 16(3), 936–938. 10.1016/j.brs.2023.05.022

Tendler, A., Roth, Y., & Harmelech, T. (2021). Deep repetitive TMS with the H7 coil is sufficient to treat comorbid MDD and OCD. Brain Stimulation, 14(3), 658–661. 10.1016/j.brs.2021.04.006

Thies, M., Zrenner, C., Ziemann, U., & Bergmann, T. O. (2018). Sensorimotor mu-alpha power is positively related to corticospinal excitability. Brain Stimulation, 11(5), 1119–1122. 10.1016/J.BRS.2018.06.006

Tonin, L., Perdikis, S., Kuzu, T. D., Pardo, J., Orset, B., Lee, K., Aach, M., Schildhauer, T. A., Martínez-Olivera, R., & Millán, J. del R. (2022). Learning to control a BMI-driven wheelchair for people with severe tetraplegia. IScience, 25(12), 105418. 10.1016/j.isci.2022.105418

Treder, M. S. (2020). MVPA-Light: A Classification and Regression Toolbox for Multi-Dimensional Data. Frontiers in Neuroscience, 14, 289. 10.3389/FNINS.2020.00289/BIBTEX

Wang, J. X., Rogers, L. M., Gross, E. Z., Ryals, A. J., Dokucu, M. E., Brandstatt, K. L., Hermiller, M. S., & Voss, J. L. (2014). Memory Enhancement: Targeted enhancement of cortical-hippocampal brain networks and associative memory. Science, 345(6200), 1054– 1057. 10.1126/SCIENCE.1252900/SUPPL_FILE/WANG-SM.PDF

Wischnewski, M., Haigh, Z. J., Shirinpour, S., Alekseichuk, I., & Opitz, A. (2022). The phase of sensorimotor mu and beta oscillations has the opposite effect on corticospinal excitability. Brain Stimulation, 15(5), 1093–1100. 10.1016/J.BRS.2022.08.005

Ziemann, U., Rothwell, J. C., & Ridding, M. C. (1996). Interaction between intracortical inhibition and facilitation in human motor cortex. Journal of Physiology, 496(3), 873–881. 10.1113/JPHYSIOL.1996.SP021734

Zrenner, C., Desideri, D., Belardinelli, P., & Ziemann, U. (2018). Real-time EEG-defined excitability states determine efficacy of TMS-induced plasticity in human motor cortex. Brain Stimulation, 11(2), 374–389. 10.1016/J.BRS.2017.11.016

