## Supplement for "Personalized whole-brain activity patterns predict human corticospinal tract activation in real-time"

**Supplementary Information for Khatri et al. 2024**

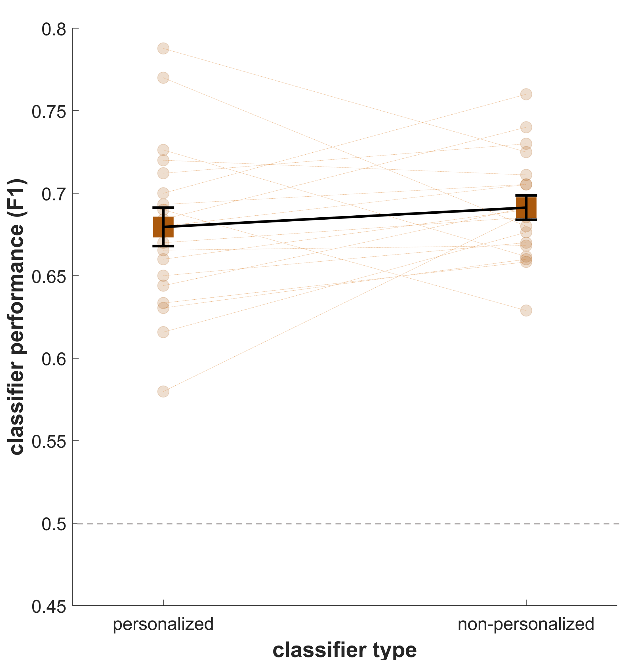

**Supplementary Figure 1. F1 values indicating performance of personalized and non-personalized classifiers.** Squares denote group averages, circles denote individual data points, error bars reflect SEM, and dashed line reflects theoretical chance level (0.5).

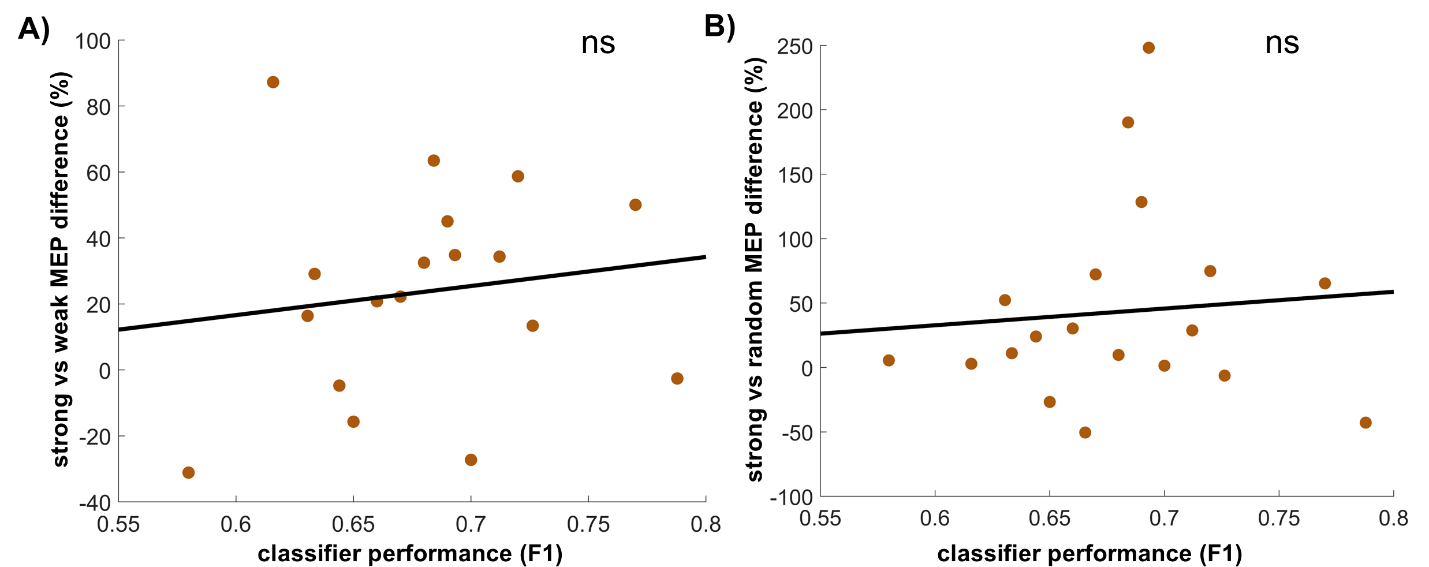

**Supplementary Figure 2. Relationships between brain state-dependent variation in CST output and personalized classifier performance.** Relationships between (A) percentage differences in MEP amplitudes elicited during strong versus weak CST states and personalized classifier F1 scores, and (B) percentage differences in MEP amplitudes elicited during strong versus random CST states and personalized classifier F1 scores. “ns” denotes lack of significance and circles denote individual subjects.

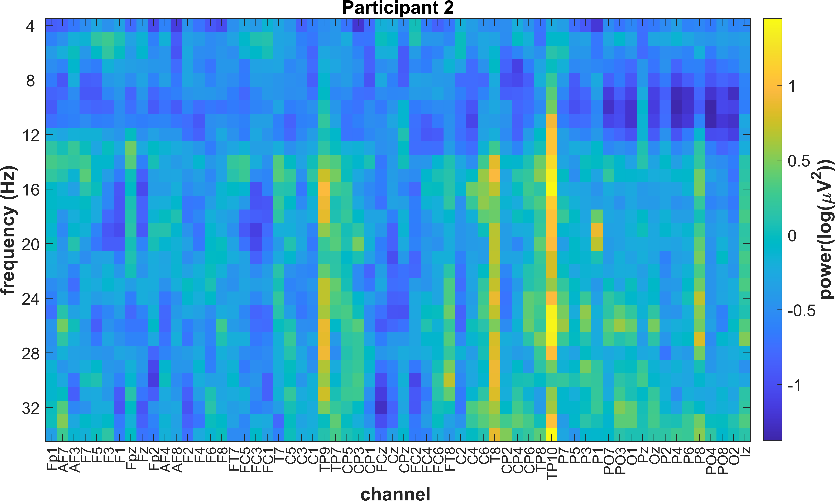

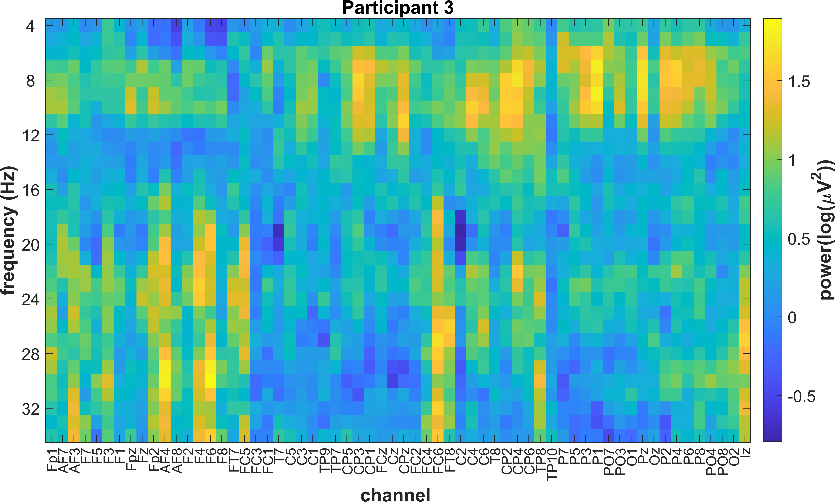

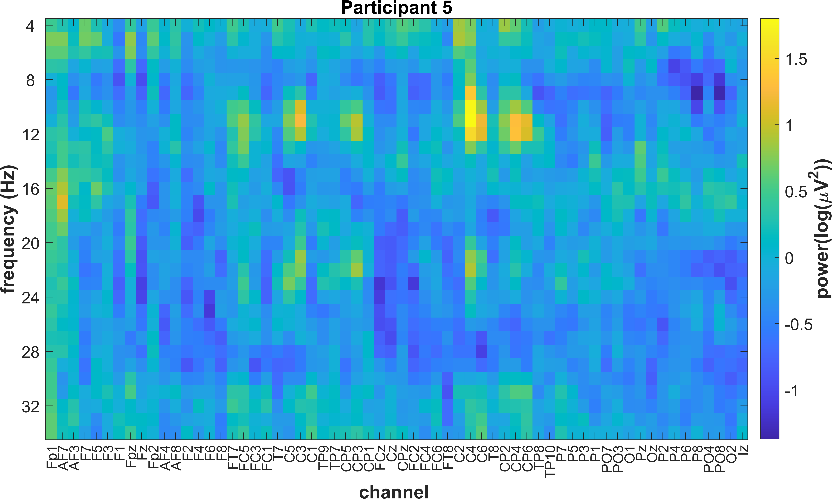

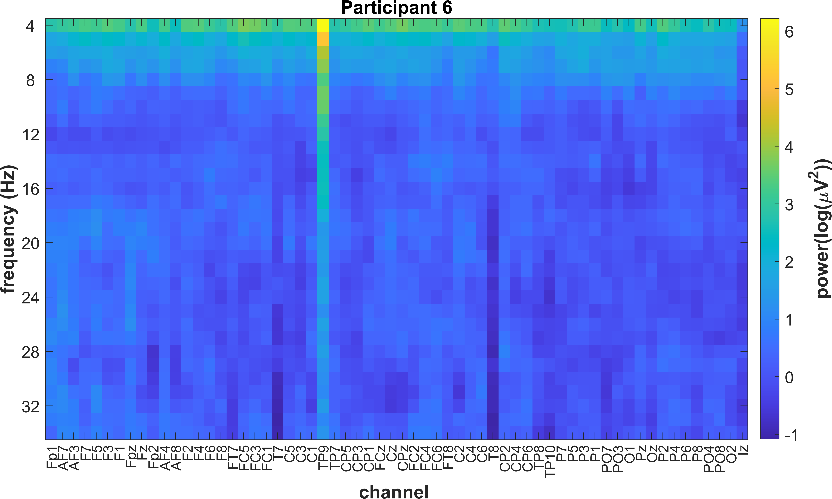

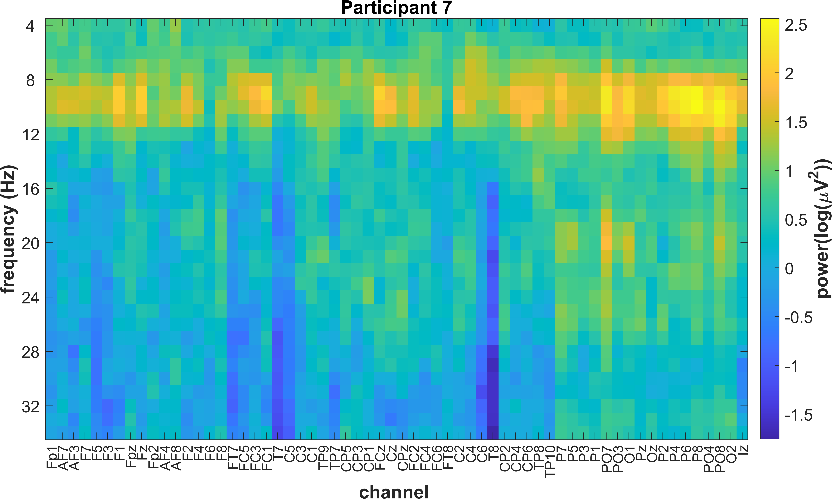

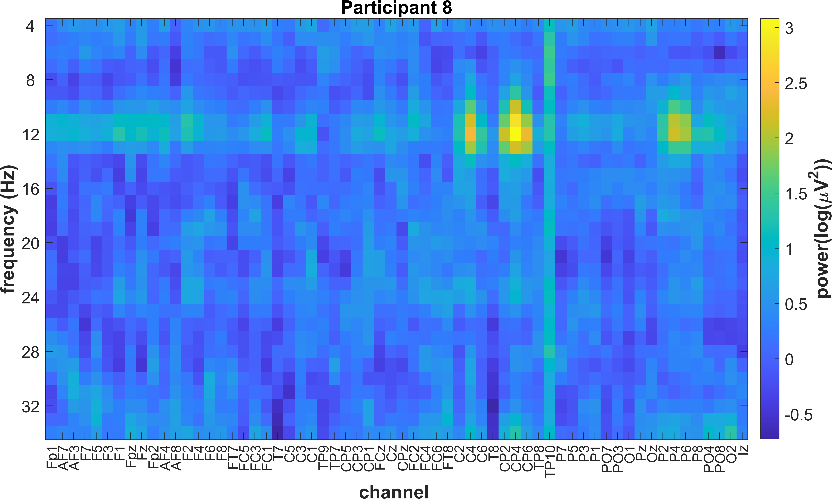

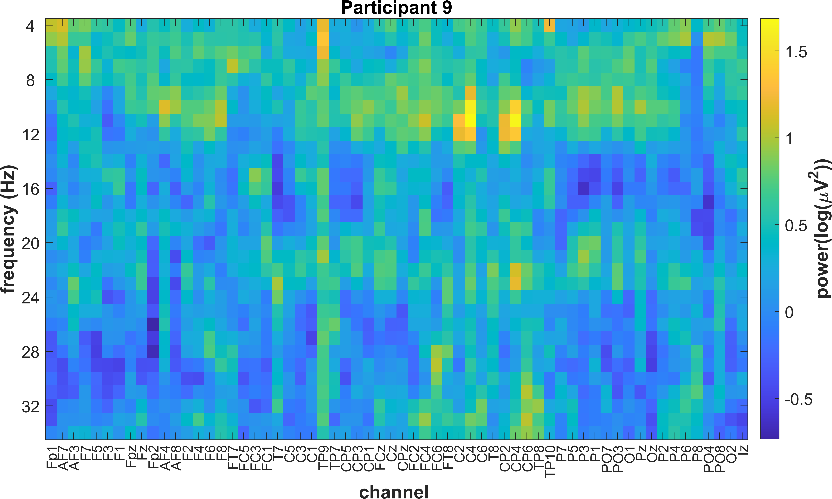

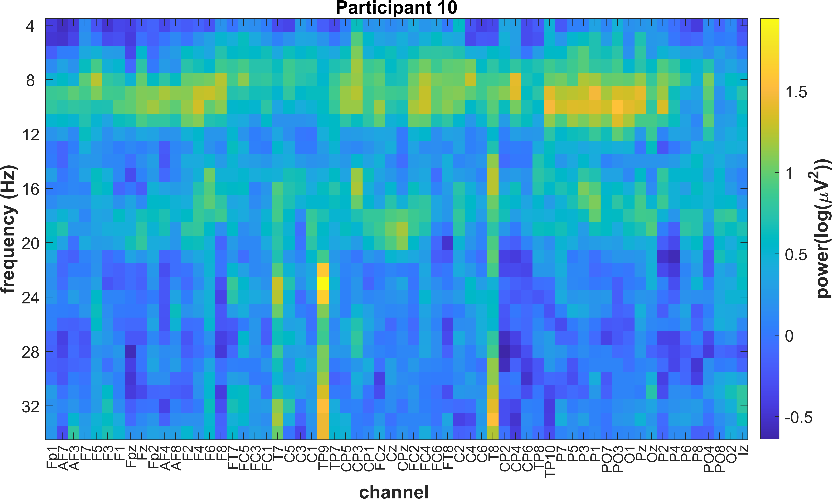

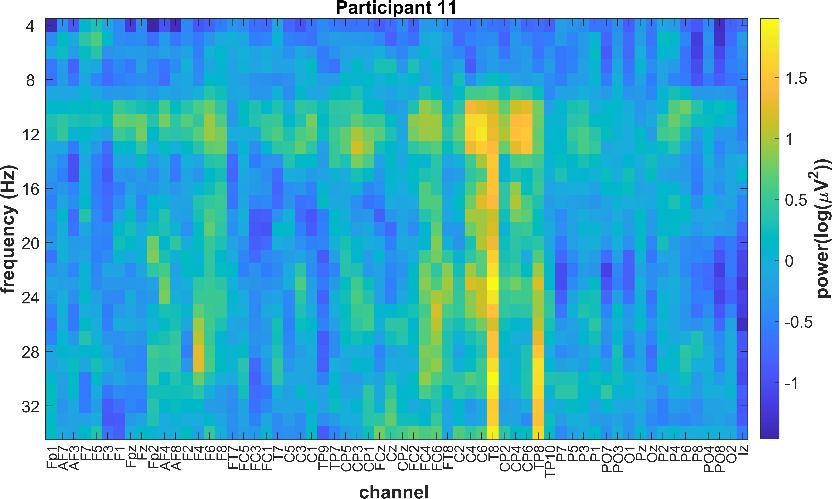

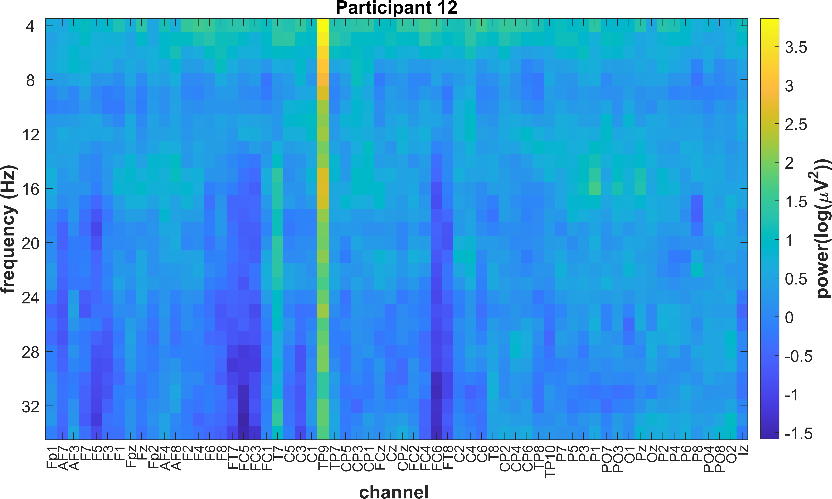

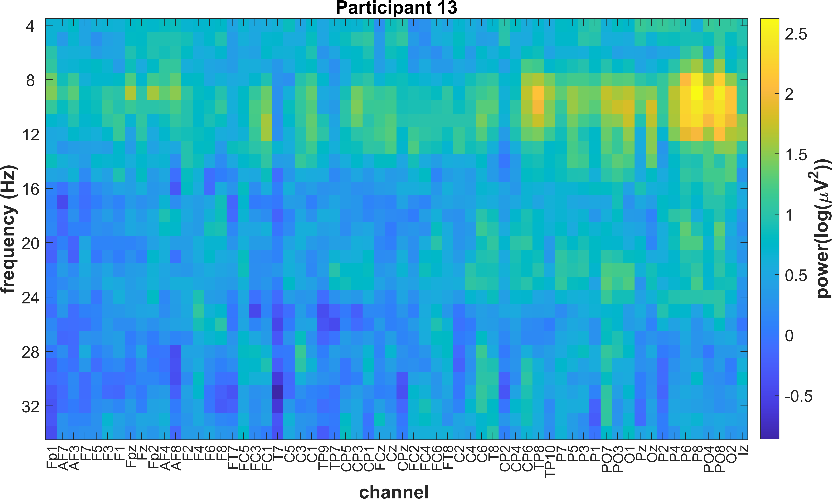

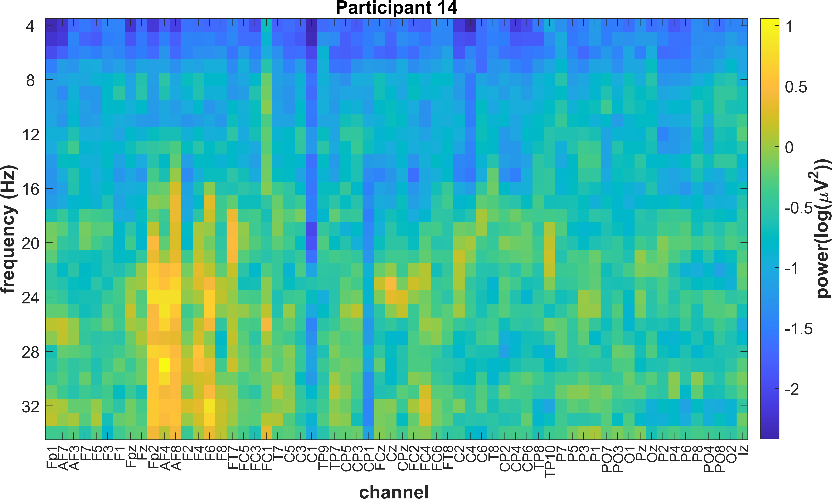

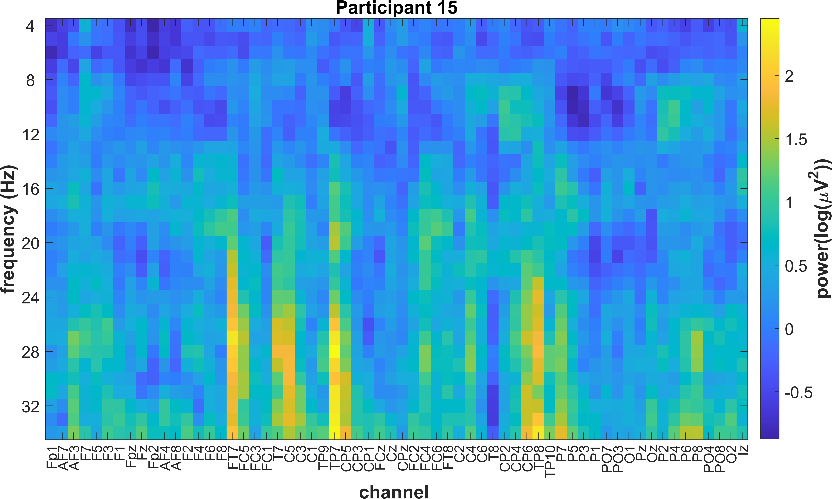

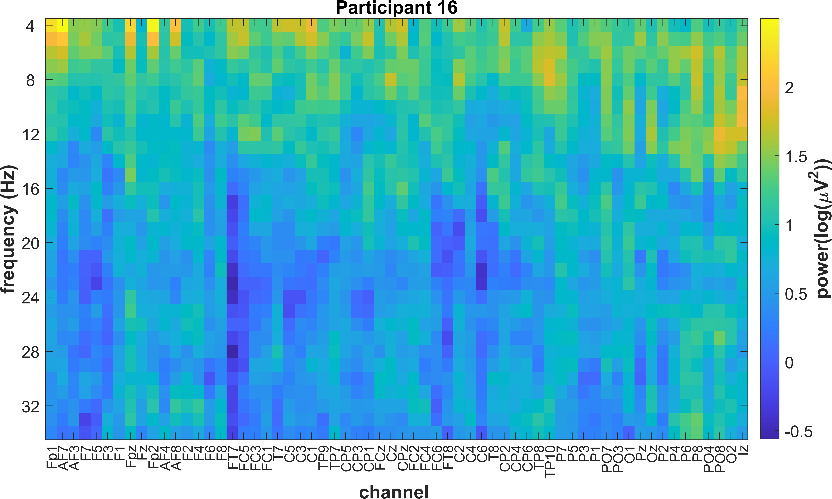

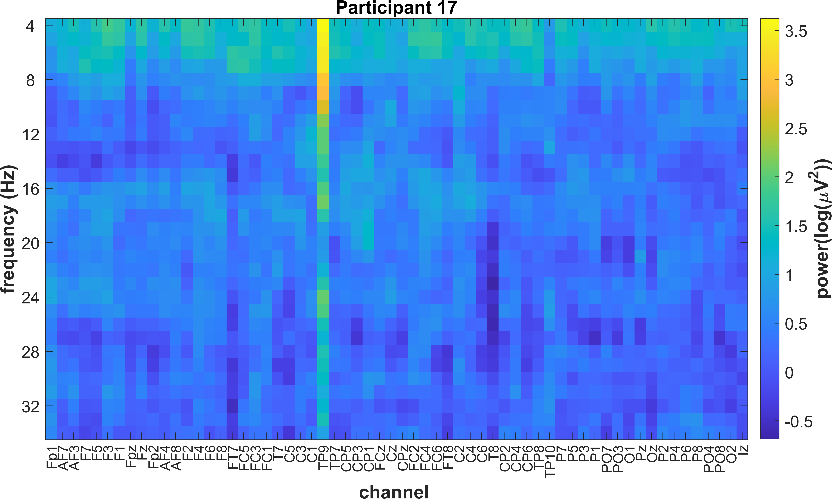

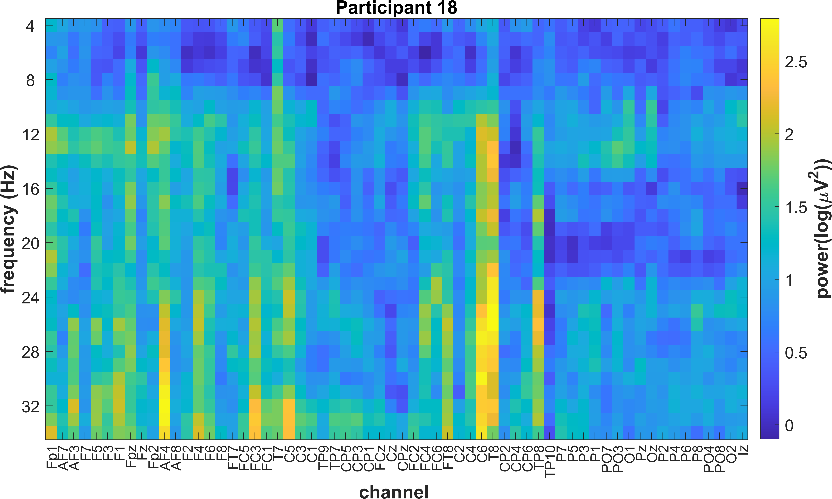

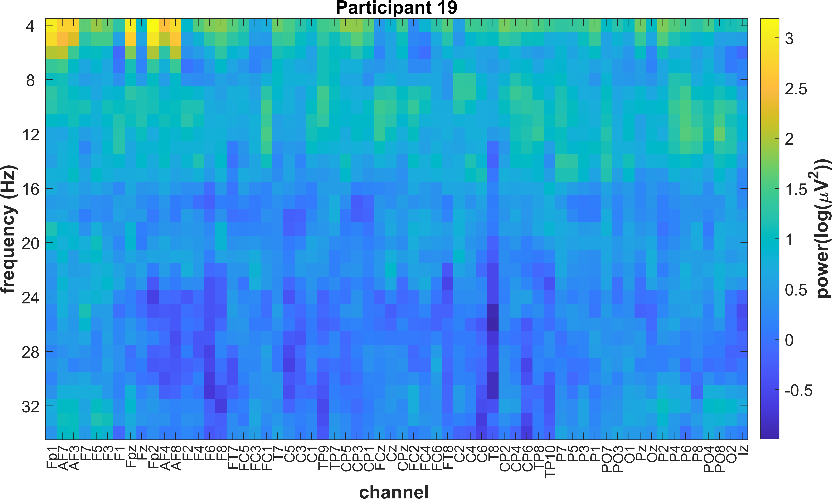

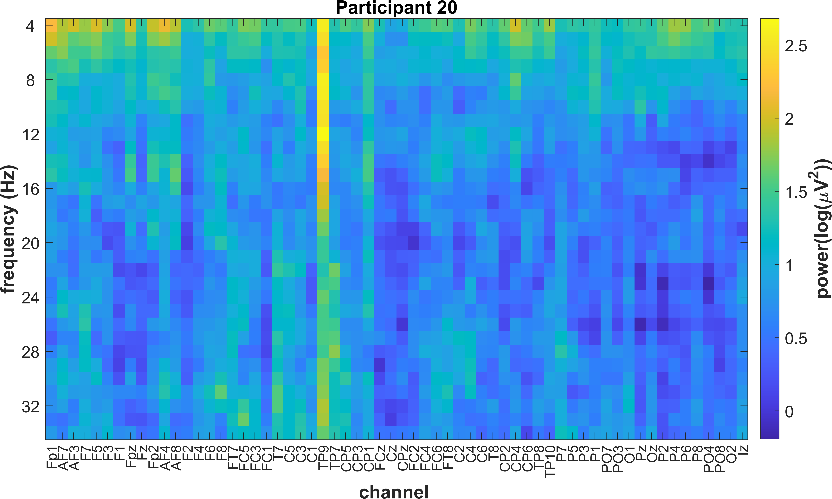

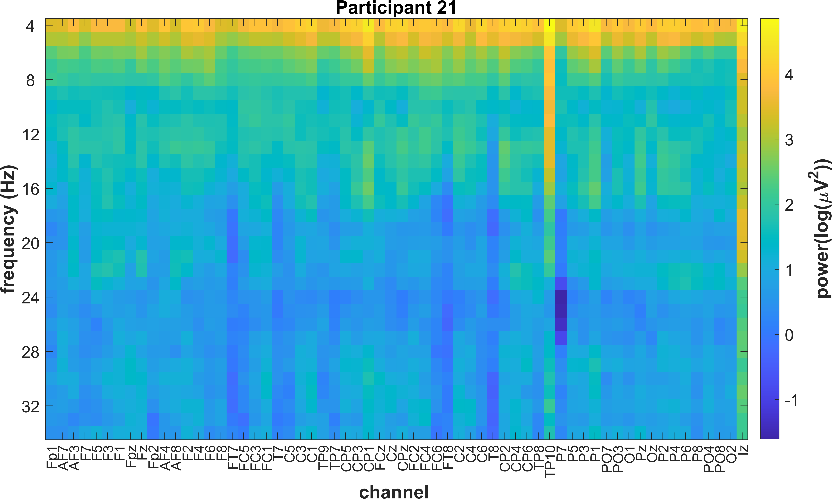

**Supplementary Figure 3. Participant-specific differences in pre-stimulus spectro-spatial EEG patterns corresponding to personalized strong and weak CST states in real-time.** For each plot, colors are scaled to individual minima and maxima to emphasize personalized EEG activity patterns.
